# Microtubule end stabilisation by cooperative oligomers of Ska and Ndc80 complexes

**DOI:** 10.1101/2025.07.06.663352

**Authors:** Renjith M. Radhakrishnan, Lauren Stokes, Matthew Day, Pim J. Huis in ’t Veld, Vladimir A. Volkov

## Abstract

During mitosis, properly aligned chromosomes stabilise microtubule ends with the help of kinetochores to ensure timely segregation of chromosomes. Microtubule-binding components of the human outer kinetochore, such as Ndc80 and Ska complexes, are present in multiple copies and together bind several microtubule ends, creating a highly multivalent binding interface. Whereas Ndc80:Ndc80 and Ndc80:microtubule binding is crucial for interface stability, Ndc80 alone in absence of Ska is unable to support stable kinetochore-attachments. Using cryoET, we demonstrate that oligomeric Ndc80:Ska assemblies stabilise microtubule ends against shortening by strengthening lateral contacts between tubulin protofilaments at microtubule plus-ends. We further identify a point mutation within the SKA1 microtubule-binding domain that does not affect microtubule-binding of individual Ska molecules, but does abolish Ska:Ska interactions. Finally, we report that oligomerisation of Ska, in a cooperative fashion together with the Ndc80, is necessary to maintain stable microtubule attachments both *in vivo* and *in vitro*.

## INTRODUCTION

During mitosis, microtubules of the mitotic spindle bind to kinetochores, large multiprotein assemblies forming at the centromeric regions of chromosomes. Having established a bioriented configuration, kinetochores mainly interact with microtubule ends (Shrestha and Draviam, 2013). These end-on attachments persist over multiple cycles of microtubule shortening and growth, supporting kinetochore motility with the filament’s end, and resisting detachment (Akiyoshi et al., 2010; Nicklas and Ward, 1994; Skibbens et al., 1993; Stumpff et al., **2008**). Strength of kinetochore’s attachment to a microtubule end is hypothesised to be important to prevent accidental chromosome loss, however an appropriate balance of strength needs to be achieved in order to correct erroneous attachments before they lead to chromosome abnormalities such as aneuploidy (Deluca et al., 2006, 2011; Liu et al., 2009; Long et al., 2017; Thompson and Compton, 2011).

The kinetochore-microtubule binding interface is highly multivalent **(Figure 1A)**. The Ndc80 complex, the main microtubule-binder conserved across a majority of eukaryotes, in human cells is present at each kinetochore in hundreds of copies interacting with 5-15 microtubule ends (Kiewisz et al., 2022; Suzuki et al., 2015). Each copy of Ndc80 can bind microtubules via two distinct regions: the globular calponin-homology domains of the NDC80 and NUF2 subunits, and the unstructured N-terminal tail of the NDC80 subunit (Ciferri et al., 2008, 2005; Wei et al., 2005) **(Figure 1B)**. The microtubule-binding regions are separated from the kinetochore-binding RWD domains of the SPC24 and SPC25 subunits by a coiled coil-rich stalk of about 58 nm.

**Figure 1.**
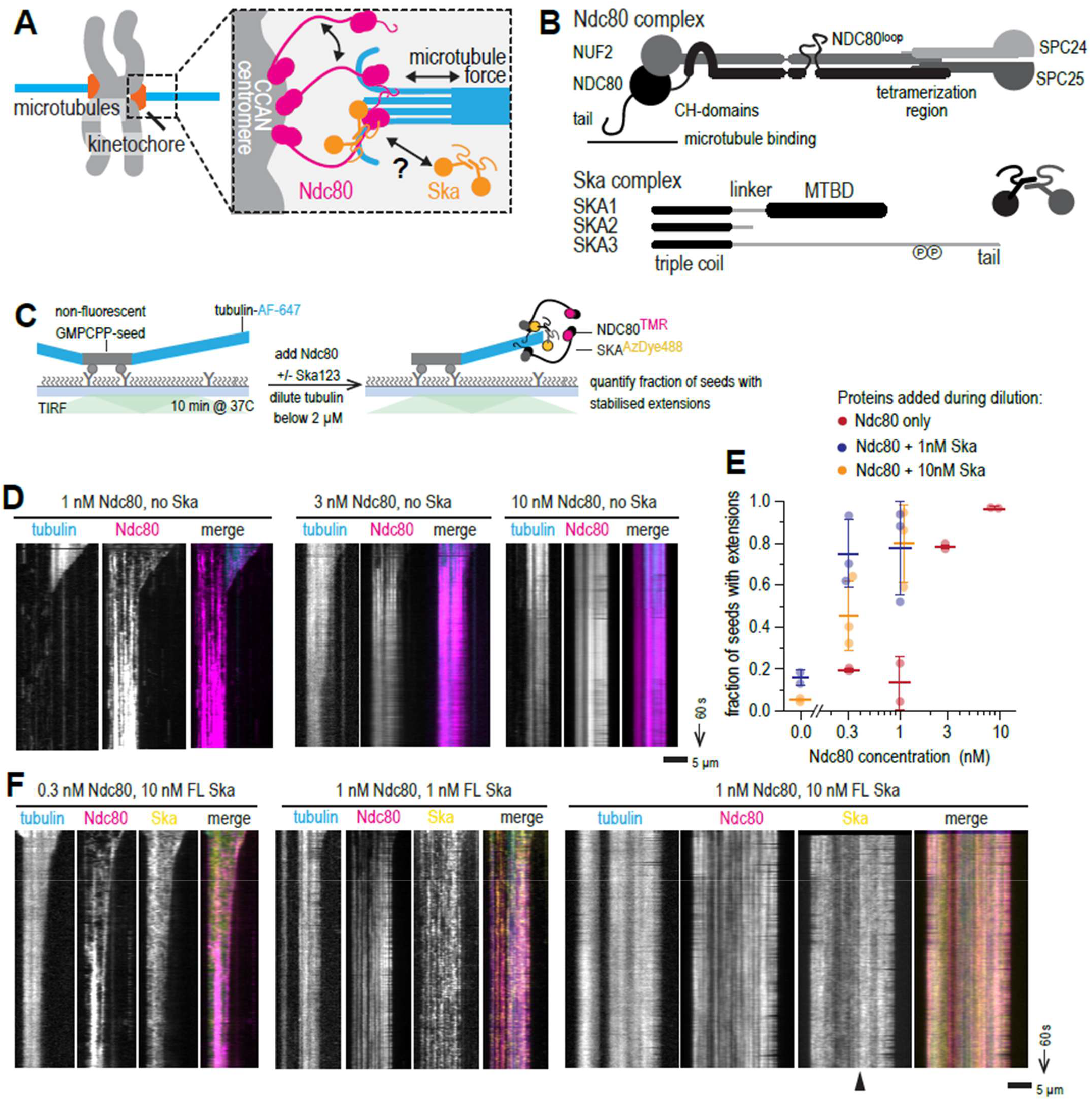
Cooperative binding of Ndc80 and Ska prevents microtubule disassembly. **(A)** Schematic of the multivalent kinetochore-microtubule interface. **(B)** Domain organisation of human Ndc80 and Ska complexes. **(C)** Schematic of an experiment to test microtubule end stabilisation by Ndc80 and Ska. **(D)** Example kymographs showing microtubule length overtime following the addition of Ndc80 at a concentration indicated. **(E)** Fraction of unlabelled GMPCPP-stabilised microtubule seeds with fluorescent tubulin extension following 10 minutes after tubulin dilution. Magenta: only Ndc80 added, blue: Ndc80 + 1nM Ska, yellow: Ndc80 + 10nM Ska (at least 50 seeds quantified in total over at least 5 fields of view per repeat, repeated 2-3 times). Lines and error bars represent mean ± SD. Two-way **AN** OVA: column factor (Ndc80 concentration) p < 0.0001; row factor (Ska concentration) p = 0.0012. **(F)** Example kymographs showing microtubule length over time following the simultaneous addition of Ndc80 and Ska at a concentration indicated. Arrowhead points to non-uniform microtubule decoration by Ska. Scale bars: 5 µm (horizontal), 60 s (vertical).

Multivalency of the Ndc80 complexes is crucial for their ability to support motility with the shortening microtubule ends *in vitro* **(Powers et al., 2009; Volkov et al., 2018)**; kinetochores multimerise Ndc80 in several ways. First, Ndc80 is recruited to the kinetochore via two complementary binding pathways: CENP C:Mis12:Ndc80 in a stoichiometry of 1:1:1 (Huis in ’t Veld et al., 2016; Petrovic et al., 2010), a complex that forms the basis for the KNL1 recruitment and checkpoint signalling (Cheeseman et al., 2006; Polley et al., 2024; Yatskevich et al., 2023), and the CENP-T:Mis12:Ndc80 pathway with a stoichiometry of 1:1 :3 in humans (Huis in ’t Veld et al., 2016; Kim and Yu, 2015; Rago et al., 2015) and 1:1:2 in chicken cells (Takenoshita et al., 2022) and budding yeast (Pekgoz Altunkaya et al., 2016). Both CENP-C- and CENP-T-dependent pathways contribute to the resulting Ndc80 copy number per kinetochore approximately equally (Suzuki et al., 2015).

Second, in conditions of high occupancy of its binding sites on the microtubule, Ndc80 CH-domains interact with their neighbours, with a proposed bridging role of the unstructured N-terminal tail in cross-linking them **(Alushin et al., 2012, 2010)**. Finally, neighbouring Ndc80s interact with each other in a cooperative manner via the loop region situated about 20 nm away from the microtubule-binding site **(Polley et al., 2023)**. We previously showed that loop-dependent Ndc80:Ndc80 binding was necessary to allow Ndc80 oligomers to rescue microtubule shortening under force, while absence of the loop region from cellular Ndc80, or point mutations in the loop, triggered the spindle assembly checkpoint (SAC) response and arrested the cells in prometaphase **(Polley et al., 2023)**. Thus, the microtubule-binding activity of the Ndc80 is insufficient for its ability to bind and stabilise microtubule ends, and needs to be supplemented with allosteric oligomerisation.

In humans and other related animals, Ndc80 recruits an additional microtubule-binding complex called Ska. Human Ska complex consists of three polypeptides: SKA1, harbouring a C-terminal microtubule-binding domain (MTBD), SKA3 with an extended C-terminal tail that is required for Cdk1-dependent Ndc80 binding, and SKA2, which stabilises N-termini or all three subunits via formation of a triple coil in the form of a winged helix domain (Abad et al., 2014; **Jeyaprakash et al., 2012; Zhang et al., 2017) (Figure 1B)**. Recruitment of the Ska complex is one of the latest mitotic events, however it is essential for proper mitotic progression, stability of kinetochore microtubule attachments, and satisfaction of SAC (Auckland et al., 2017; Cheerambathur et al., 2017; Daum et al., 2009; Gaitanos et al., 2009; Hanisch et al., 2006; Raaijmakers et al., 2009; Sivakumar et al., 2016). While it is currently unclear how many copies of Ska can be recruited at the kinetochore, *in vitro* it binds Ndc80 in a stoichiometric manner (Huis in ’t Veld et al., 2019) and thus is likely to be oligomerised, at least via the pathways described above leading to Ndc80 oligomerisation by its inner kinetochore receptors. Previous studies reporting Ska:Ndc80 interactions *in vitro* used either incomplete protein fragments **(Schmidt et al., 2012)**, or full complexes either in homogenous micromolar mixtures **(Huis in ’t Veld et al., 2019)**, or oligomerised by beads for optical trapping (Helgeson et al., 2018; Huis in ’t Veld et al., 2019). Thus it is currently unclear whether full-length Ska and Ndc80 can spontaneously oligomerise on microtubules, or they require externally imposed oligomerisation.

*In vitro*, purified Ska was shown to follow both growing and shortening microtubule ends (Auckland et al., 2017; Maciejowski et al., 2017; Monda et al., 2017; Welburn et al., 2009), and to enhance end tracking ability of Ndc80 with impaired intrinsic end tracking: either non-end-tracking fragments (Schmidt et al., 2012), or tailless but otherwise full-length oligomers of Ndc80 (Huis in ’t Veld et al., 2019). Microtubule end tracking was attributed to the affinity of Ska to bent tubulin oligomers (Monda et al., 2017; Schmidt et al., 2012), conformations that are present at both growing and shortening microtubule ends (McIntosh et al., 2018). SKA1 MTBD was shown to carry multiple positively charged residues that contributed to its interaction with either growing or shortening microtubule ends, and to proper mitotic timing (Abad et al., 2014; Monda et al., 2017).

Despite the almost ubiquitous conservation of the Ndc80 complex in eukaryotes, its binding partners are divergent: while most animals have some versions of the Ska complex, Fungi lack it and instead cross-link their Ndc80 to microtubule ends with a ring-forming Dam1 complex **(Miranda et al., 2005; van Hooff et al., 2017; Westermann et al., 2005)**. Dam1 and Ska share no sequence or domain conservation, however they both are essential for cell viability and contribute to the mitotic fidelity through the same general mechanism: cross-linking the coiled coil of the Ndc80 complex to the tubulin flare at the end of a dynamic microtubule (Helgeson et al., 2018; Huis in ’t Veld et al., 2019; Lampert et al., 2010; Muir et al., 2023; Schmidt et al., 2012; Tien et al., 2010; Wimbish et al., 2020). Dam1 complex can assemble into ring-shaped oligomers thanks to interactions between its heterodecameric subunits (Jenni and Harrison, 2018; Miranda et al., 2005; Muir et al., 2023; Westermann et al., 2005), assisted by their binding to the Ndc80 complex, Bim1^EB1^, and microtubules (Dudziak et al., 2021; Muir et al., 2023; Westermann et al., 2005). Mutations affecting Dam1 oligomerisation lead to a reduction in kinetochore-microtubule attachment strength, and severe mitotic phenotypes (Dudziak et al., 2021; Muir et al., 2023).

While Ska complex has not been shown to form microtubule-encircling rings, its functional similarity to the Dam1 complex has provided reasons to hypothesise that it should oligomerise as well (**Jeyaprakash et al., 2012; Maciejowski et al., 2017; Welburn et al., 2009**). Indeed, chemical cross-linking of the soluble Ska complex followed by mass-spectrometry have identified some regions within each of the three Ska subunits whose proximity might be sufficient for a direct interaction (Helgeson et al., 2018; Huis in ’t Veld et al., 2019). However, it is unclear whether any of the previously identified molecular interfaces target specifically Ska:microtubule, or Ska:Ska interactions, or their combination.

In summary, the Ska:Ndc80:microtubule system is highly cooperative, with each component contributing to stabilising the other components: microtubules via their regular lattice structure, Ndc80 and Ska via their interactions with themselves, and with each other. The following properties appear to be essential but not individually sufficient for proper, stable kinetochore microtubule attachments: microtubule-binding of Ndc80, cooperative oligomerisation of Ndc80, microtubule-binding of Ska, Ska:Ndc80 interaction **(Figure 1A)**. Here we add one previously missing item to this list: we report that Ska is oligomerising via at least two interfaces; oligomerisation of SKA1 MTBD can be disrupted by a point mutation and, while not necessary for individual molecules’ microtubule binding, is indeed necessary to stabilise microtubule ends against disassembly both *in vitro* and *in vivo*.

## RESULTS

### Cooperative binding of Ndc80 and Ska prevents microtubule disassembly

In order to dissect the specific effects of Ska:Ska and Ndc80:Ndc80 interactions within the outer kinetochore system, we sought to reconstitute the ability of Ska:Ndc80 oligomers to stabilise microtubule ends *in vitro* in a minimal system of purified components. To this end, we purified full-length Ndc80 and Ska complexes, labelled them fluorescently **(Supplementary Figure 1A)**, and relied on their ability to form homo- and hetero oligomers to stabilise microtubule ends. Microtubules were grown in a flow chamber from coverslip-attached stable seeds using fluorescently labelled tubulin, and then the solution was rapidly changed to the one containing low concentrations of either Ndc80, or Ska, or both, and tubulin in a concentration below 2 µM, to induce microtubule shortening **(Figure 1C)**. Ndc80 alone was able to stabilise microtubule ends in concentrations above 3 nM; Ndc80 concentrations of 1 nM and below resulted in microtubule depolymerisation **(Figure 1D)**. To quantify the observed stabilising effect, we counted the fraction of seeds that carried fluorescent tubulin extensions after 10 minutes post-dilution, and found a sharp transition in the amount of stabilised seeds between 1 and 3 nM of added Ndc80 **(Figure 1E)**, consistent with the previously reported cooperative recruitment of Ndc80 to microtubules in the same concentration range (Polley et al., 2023).

Ska alone was unable to stabilise microtubules against shortening at concentrations as high as 10 nM **(Figure 1 E)**. However, presence of 1-10 nM Ska in addition to low, non-stabilising amounts of Ndc80, resulted in stabilisation of microtubule ends and recruitment of Ska and Ndc80 oligomers to microtubule ends and lattices **(Figure 1EF)**. At 10 nM Ska, we also observed higher density decoration of Ska towards the stabilised microtubule plus-end (**Figure 1F**, arrowhead), which we will discuss in detail below. Two-way ANOVA confirmed that both the concentration of Ndc80, and the concentration of Ska contributed significantly to an increase in the fraction of fluorescent tubulin extensions (p < 0.0001 for the effect of the Ndc80 concentration; p = 0.0006 for the effect of Ska concentration).

Phosphorylation of SKA3 at T358 and T360 was previously shown to be essential for Ska:Ndc80 interaction in vitro (Huis in ’t Veld et al., 2019) and kinetochore recruitment of Ska (Zhang et al., 2017), however we previously demonstrated that Ska expressed in insect cells and purified without further modifications was partially phosphorylated allowing it to interact with Ndc80, unless specifically dephosphorylated using a phosphatase (Huis in ’t Veld et al., 2019). To test whether the Ska:Ndc80 interaction in our microtubule end-stabilisation assay required additional phosphorylation on top of the background phosphorylation introduced during insect cell expression, we used purified Cdk1: CyclinB:CKS1 complex to phosphorylate purified FL Ska in vitro, resulting in a prominent shift in SKA3 migration on SDS-PAGE (**Supplementary Figure 1B**). Repeating the microtubule end-stabilisation assay in presence of hyperphosphorylated or untreated FL Ska, we did not observe a statistically significant difference in the fraction of seeds carrying fluorescent microtubule extensions (2-way ANOVA: p = 0.0003 for the effect of the Ndc80; p = 0.33 for the effect of Ska hyperphosphorylation, **Supplementary Figure 1C**). In addition, both hyperphosphorylated and untreated Ska molecules were co-localising with Ndc80 on microtubule lattice (**Supplementary Figure 1D**). Based on these observations, we pooled results with both hyperphosphorylated and untreated Ska.

### Ska:Ndc80 oligomers stabilise microtubule plus-ends by reinforcing lateral interactions between bent protofilaments

In order to gain a mechanistic understanding of microtubule end stabilisation by Ska and/or Ndc80 oligomers, were repeated the tubulin dilution experiments on grids suitable for cryoET. Microtubules were grown from stable seeds attached to the silanized and passivated holey SiO support film, and then diluted with Ska and/or Ndc80 containing solution and incubated for 2 min before plunge-freezing (**Figure 2A**). Examining the tomograms containing microtubule plus-ends in presence of the stabilising concentrations of Ndc80 alone, we observed oligomers of Ndc80, often forming long “trains” with their CH-domains and apparently intertwined with their C-terminal coiled coils, bound near the microtubule ends and along the lattices **(Figure 2B, Supplementary Figure 2A)**. With 10 nM Ska present in addition to 1nM Ndc80, we observed similar oligomers of Ndc80 **(Figure 2C)**, and, additionally, microtubule-bound oligomers that lacked the characteristic ordered “trains” of Ndc80 CH-domains with the elongated coiled-coils **(Supplementary Figure 2BC)**. Although the resolution in individual denoised tomograms was not sufficient to confidently determine structural properties within these oligomers, we attribute these additional densities to the presence of Ska.

**Figure 2.**
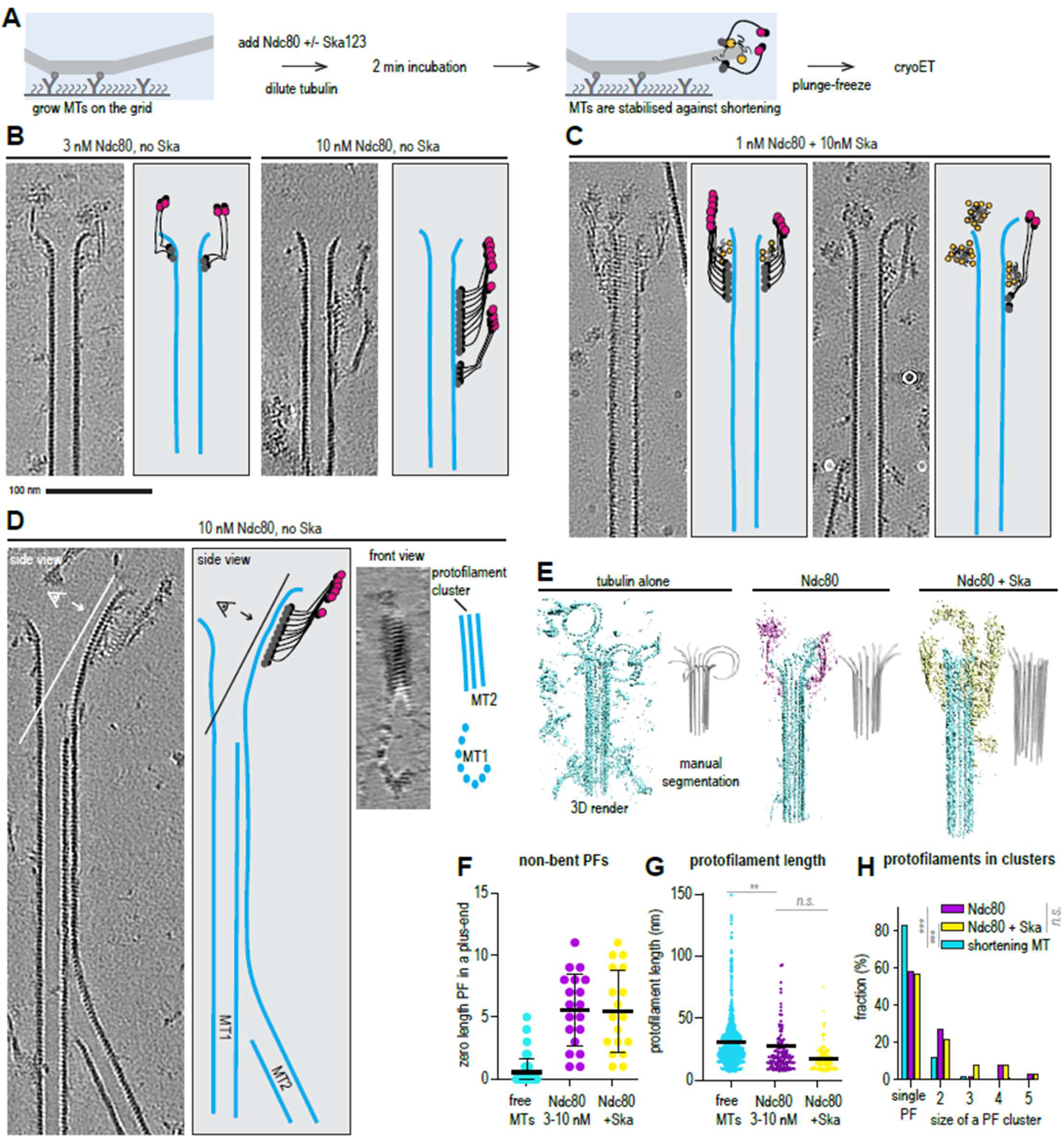
Ndc80 and Ska stabilise microtubule ends by facilitating formation of protofilament clusters. **(A)** Schematic of an experiment to test microtubule end stabilisation by Ndc80 and Ska using cryoET. **(B)** Representative 0.8-nm thick slices through cryoCARE-denoised tomograms showing microtubule end- and wall-bound oligomers of Ndc80 (greyscale), and their interpretation (colour) side by side. **(C)** Representative tomographic slices showing microtubule end- and wall-bound oligomers in presence of Ndc80 and Ska. **(D)** Example of an Ndc80 oligomer stabilising an extended, sheet-like protofilament cluster, viewed from the side (left), or from the luminal side of the cluster (right). **(E)** Examples of rendered 3D densities containing a microtubule plus-end, and the result of its manual segmentation to obtain the shapes of protofilaments for three conditions: undecorated shortening microtubules (n = 65 MT ends, 888 protofilaments, previously published dataset available at EMPIAR-12554), and microtubules stabilised form shortening using Ndc80 (n = 19 MT ends, 150 protofilaments) or Ndc80 + Ska (n = 18 MT ends, 144 protofilaments). **(F)** Amount of protofilaments without a bent part in a microtubule plus-end. **(G)** Protofilament length at microtubule plus-ends stabilised by Ndc80 or Ndc80 + Ska. Protofilament lengths for free shortening microtubule ends are plotted from a previously published dataset (Kalutskii et al., 2025). Welch’s t-test p-values: Ndc80 vs free shortening: 0.4089; Ndc80 vs Ndc80 + Ska: 0.0036; Ndc80 + Ska vs free shortening: <0.0001. **(H)** Fraction of protofilaments engaged in laterally associated clusters, sorted by cluster size. Chi-squared test p-value: Ndc80 vs Ndc80 + Ska: 0.1788; Ndc80 vs free shortening:< 0.0001; Ndc80 + Ska vs free shortening: < 0.0001. Scale bars: 100 nm.

To eliminate the possibility that we observed the ends of stable GMPCPP seeds, instead of GDP ends prevented from shortening by Ndc80 and Ndc80+Ska, we performed additional checks. First, we analysed the ends of those microtubules that were longer than the typical length of GMPCPP-seeds, as determined by TIRF microscopy **(Supplementary Figure 2D)**. Second, building on previous observations that GMPCPP stabilised microtubules have expanded lattices compared to GDP-bound microtubules **(Manka and Moores, 2018; Zhang et al., 2015)**, we correlated the microtubule length to the tubulin lattice spacing near the plus ends and excluded plus ends of short expanded microtubule seeds from further analysis **(Supplementary Figure 2E)**. Finally, we compared the lattice spacing of the selected plus ends to our previously published dataset containing shortening microtubule ends in absence of decoration by other proteins and found a similar compacted spacing of (Kalutskii et al., 2025) **(Supplementary Figure 2F)**.

How do Ndc80 and Ndc80/Ska oligomers prevent microtubules from shortening? We found a possible clue to this question by observing extended sheet-like protofilament structures stabilised at their plus-ends by Ndc80 oligomers **(Figure 2D, Supplementary Figure 2C)**. In these structures, several protofilaments appeared to be cross-linked laterally. We previously reported that increased lateral clustering of tubulin protofilaments is a property that is characteristic of the GTP-bound, or growing microtubule ends (Kalutskii et al., 2025). We therefore set out to test a hypothesis that Ndc80 and Ndc80/Ska oligomersprevent microtubule shortening by strengthening lateral interactions between tubulin protofilaments at microtubule plus-ends. To test it, we manually traced the 3D shapes of protofilaments in 19 microtubule plus ends in each condition **(Figure 2E)**, and compared them to microtubules shortening in absence of stabilising proteins **(Kalutskii et al., 2025)**. We found that in both stabilising conditions, 3-10 nM Ndc80 alone and **1** nM Ndc80 + 10 nM Ska, microtubule plus-ends carried many more protofilaments without any bent segments, a conformation rarely observed with free dynamic microtubules **(Figure 2F)**. Despite this difference, the length of bent protofilament segments of Ndc80-stabilised microtubules was not significantly different from free shortening microtubule plus-ends **(Figure 2G**, Welch’s t-test p = 0.4). On the other hand, microtubule plus-ends decorated with Ndc80 in presence of Ska did carry shorter bent protofilaments than Ndc80 alone, or than free ends **(Figure 2G**, p = 0.0036 for Ndc80 with and without Ska).

We further tested if these bent protofilaments were interacting laterally with each other. We found that both Ndc80 alone, and Ndc80 with Ska, induced clustering of 42.0% and 42.4% of protofilaments, respectively, twice or more the amount observed at free plus ends (16-22% **(Kalutskii et al., 2025))**. Chi-squared analysis showed similar distributions of the number of protofilaments belonging to each cluster size for both Ndc80 alone, and Ndc80 with Ska **(Figure 2H**, p = 0.17), but a significant difference between each of these datasets, and freely shortening microtubule plus-ends **(Figure 2H)**. Taken together, results presented above indicate that Ndc80 oligomers stabilise microtubule ends by reinforcing lateral contacts between tubulin protofilements, and Ska acts by lowering the Ndc80 concentration that is required to achieve this effect.

### Ska oligomerises on dynamic microtubules

We further performed sub-tomogram averaging to gain a better understanding of the mechanisms that allow Ndc80 and Ska oligomers to stabilise lateral contacts between tubulin protofilaments. Focusing on microtubule-bound “trains” of Ndc80 CH-domains, we obtained low-resolution sub-tomogram averages in two separate samples described above, 3 nM and 10 nM of Ndc80 **(Figure 3A)**. We also performed the same approach on “trains” of Ndc80 CH-domains in the sample containing 1 nM Ndc80 and 10 nM Ska **(Figure 3B)**. In all three samples, we observed Ndc80 CH domains occupying two neighbouring protofilaments within the microtubule wall. However, when we repeated the same subtomogram averaging approach, picking particles from non-Ndc80 microtubule decoration in the sample containing both Ndc80 and Ska **(Figure 3B, Supplementary Figure 2B)**, we did not observe any regular structures along the microtubule lattice in our low-resolution reconstructions. Repeated rounds of 3D classification with these particles failed to yield separate classes.

**Figure 3.**
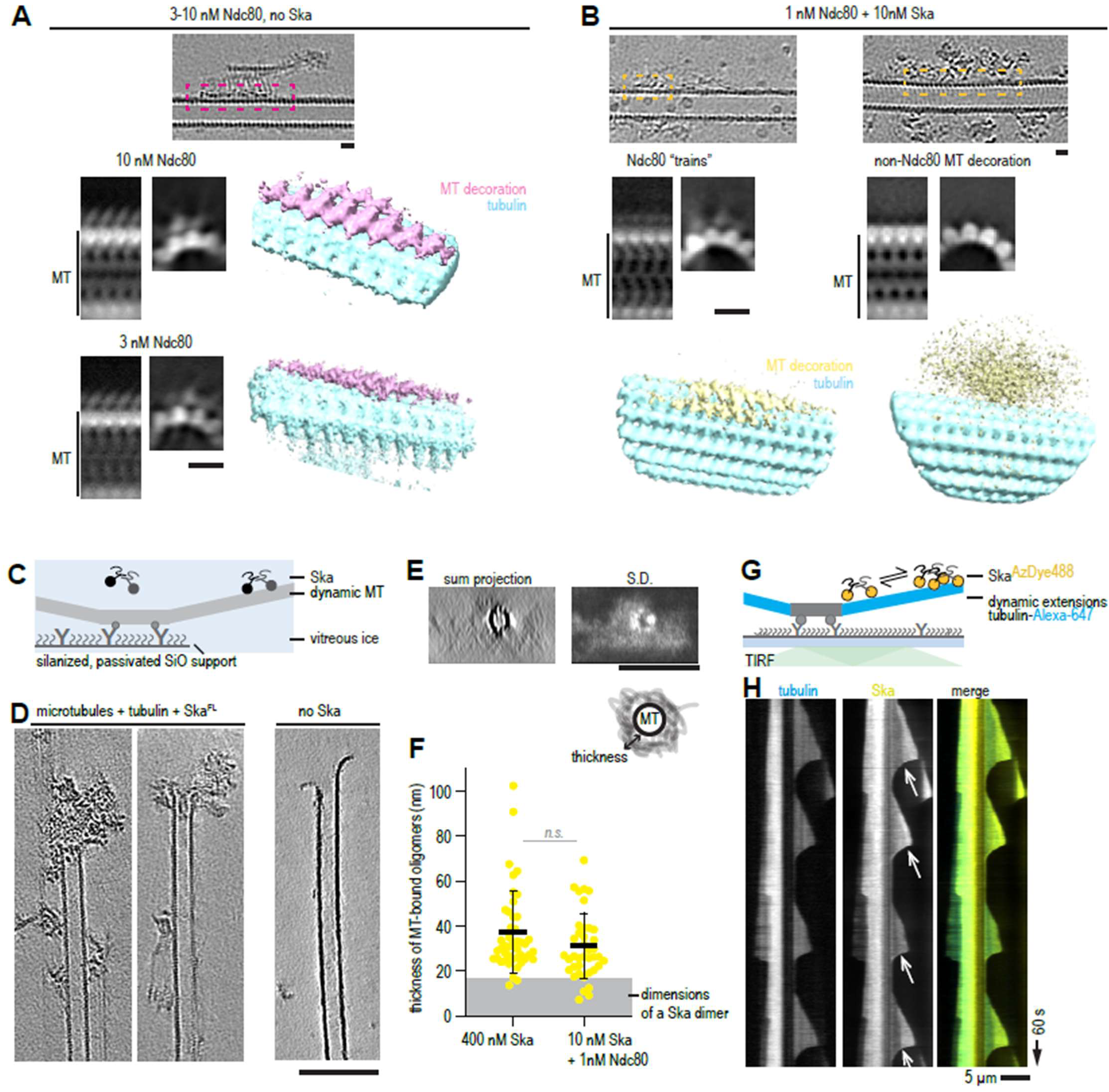
Ndc80 and Ska oligomerisation on dynamic microtubules. **(A)** Subtomogram averaging of Ndc80 CH-domain trains. White-on-black images represent 2D projections of 3D classes. Colour images show 3D rendering of the same classes. 10 nM Ndc80: 5766 particles obtained from 43 tomograms. 3 nM Ndc80: 1285 particles obtained from 26 tomograms. **(B)** Subtomogram averaging of CH-domain trains (left), and non-Ndc80 microtubule decorations in the sample containing 1 nM Ndc80 and 10 nM Ska. White-on-black images represent 2D projections of 3D classes. Colour images show 3D rendering of the same classes. Ndc80 “trains”: 822 particles obtained from 42 tomograms. Non-Ndc80 densities: 5026 particles from the same 42 tomograms. Scale bars: 10 nm. **(C)** Experimental setup to reconstitute the interaction of Ska without Ndc80 on dynamic microtubules for cryoET. **(D)** Examples of 0.8-nm thick slices through cryoCARE-denoised tomograms showing microtubule ends in presence (left) or absence (right) of Ska. Scale bar: 100 nm. **(E)** Axial projections (sum, left; and standard deviation, right) along microtubules in presence of Ska, along with a schematic diagram showing how the thickness of Ska decoration was measured. **(F)** Distribution of thicknesses of Ska decorations measured in tomograms. Yellow circles show individual measurements, lines show mean ±SD.Welch’s t-test p value: 0.0989 (n.s.). N = 45 (Ska), 39 (Ska+ Ndc80). Grey shaded area represents the expected dimensions of a Ska dimer (18 nm). **(G)** Schematic ofthe experimental setup to reconstitute microtubule binding of FL Ska at 100 nM using TIRF imaging. **(H)** A kymograph showing non-uniform decoration of microtubules (cyan) by FL Ska (yellow). White arrows show the persistent boundary between a brighter Ska envelope proximal to the growing microtubule end, and the rest of less brightly decorated microtubule lattice. Scale bar: 5 µm (horizontal), 60 s (vertical).

To get a better understanding of the nature of the Ska coating on microtubules, we repeated the sample preparation with dynamic microtubules in the presence of Ska, but without adding Ndc80 **(Figure 3C)**. We observed multiple tubulin rings cross-linked to microtubules and to each other **(Supplementary Figure 3A)**, in agreement with a previous report (Monda et al., 2017). We also observed large microtubule end-bound oligomers which we interpreted as microtubule end tracking Ska, as they were not present in absence of Ska **(Figure 3D)**. Consistently, oligomers with similar dimensions were also found in samples in which Ndc80 (1nM) and Ska (10 nM) were bot present **(Figure 3B, Supplementary Figure 2B)**. Using a previously established method of measuring overall dimensions of these oligomers extending in the direction perpendicular to the surface of the microtubule cylinder **(Maan et al., 2023)**, we found that the majority of them exceeded the expected dimensions of a single Ska dimer (Jeyaprakash et al., 2012) **(Figure 3F)**.

When we repeated the experiment in a flow chamber suitable for TIRF microscopy **(Figure 3G)**, we observed that microtubule-bound Ska formed brighter regions near the growing microtubule ends, separated from the rest of Ska-decorated microtubule lattice by a clear stationary boundary that persisted until catastrophe **(Figure 3H)**. Inspired by an analogous observations made with the microtubule-binding tau, we called these brighter regions “envelopes” **(Siahaan et al., 2019)**. Interpreting these results together with the results of our cryoET analysis, we hypothesised that non uniform decoration of microtubules by Ska is a property arising from Ska:Ska interactions, which we dissect in detail below.

### Formation of Ska envelopes requires a specific interaction of dimeric SKA1 MTBDs

To identify domains of the Ska complex that mediate Ska:Ska interactions, we mixed a crowding agent (PEG) with purified Ska carrying deletions of either the SKA3 C terminus, the SKA1 MTBD, or both **(Supplementary Figure 3B)**. Observing the resulting mixtures using fluorescence microscopy, we found round or irregular aggregates or droplets in most of the cases **(Supplementary Figure 3B, C)**. Ska^SKA3ΔC^ was the only construct displaying Ska:Ska interactions in absence of any additional crowding, however the resulting structures were not spherical **(Supplementary Figure 3B, C)**. The only Ska construct that failed to form any self interacting assemblies was Ska^SKA1ΔMTBDSKA3Δ C^ that only retained the triple coil domains **(Supplementary Figure 3D)**. Thus, Ska can interact with itself via multiple regions.

We then repeated the envelope formation assay using the tailless Ska^SKA3ΔC^, **(Figure 4A, B}**, however we found out that envelope formation was still present **(Figure 4C)**. We then probed the stability of these envelopes, and of their boundaries with the less densely decorated microtubule lattice, by recording fluorescence recovery after photobleaching (FRAP). We found that both FL Ska and Ska^SKA3ΔC^ supported formation of envelopes that exchanged with the soluble pool of Ska, and retained the intact boundary after FRAP **(Figure 4D)**. However, while Ska^SKA3ΔC^ envelopes recovered fully to pre-bleach levels, FL Ska envelopes only recovered about 60% of their initial fluorescence **(Figure 4D)**. We thus conclude that envelope formation is likely mediated by an interaction between SKA1 MTBDs, but envelopes that have formed are then stabilised by SKA3 C-terminus.

**Figure 4.**
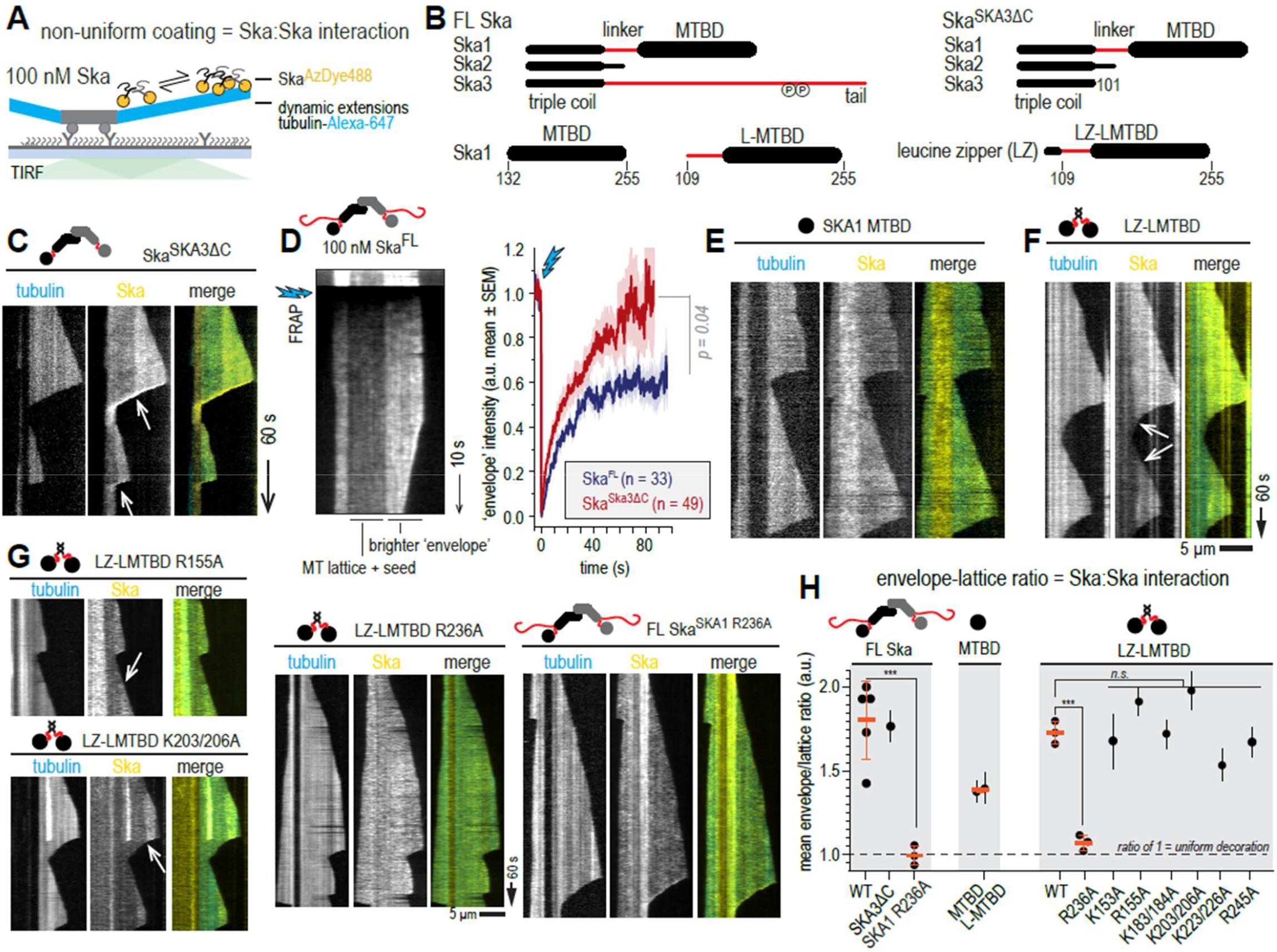
Formation of Ska envelopes requires a specific interaction of dimeric SKA1 MTBDs. **(A)** Schematic of the assay to assess Ska:Ska interaction on microtubules using its ability to form brighter envelopes as a readout. **(B)** Schematic representations of Ska constructs used in the study. Black represents folded domains, red represents disordered regions. **(C)** A kymograph showing envelope formation by Ska^SKA3ΔC^ White arrows show the persistent boundary between a brighter Ska envelope proximal to the growing microtubule end, and the rest of less brightly decorated microtubule lattice. **(D)** Example of a kymograph showing a FRAP experiment with a microtubule decorated with FL Ska (left). The graph on the right shows average FRAP curves for FL Ska and Ska^SKA3ΔC^ **(E-F)** Monomeric SKA1 MTBD **(E)** does not form a clear boundary between a brighter envelope and the rest of the microtubule lattice, as opposed to the dimeric LZ-LMTBD construct **(F). (G)** Mutants of LZ-LMTBD and FL Ska ^SKA1 R236A^ decorating dynamic microtubules. **(H)** Quantification of the extent of the envelope formation, showing mean envelope/lattice ratio of fluorescence intensity± SEM (black symbols and error bars, at least 20 microtubules per condition). Orange lines show mean ± SD of repeated experiments. Welch’s t-test: FL Ska WT vs FL Ska^SKA1 R236A^ p = 0.0009; LZ-LMTBD WT vs LZ-LMTBD R236A p = 0.0003; FL vs LZ-LMTBD p = 0.53; FL Ska^SKA1 R236A^ vs LZ-LMTBD R236A p = 0.158; LZ-LMTBD WT vs non-R236A mutants p = 0.79. Scale bar: 5 µm (horizontal), 60 s (vertical).

Surprisingly, when we repeated this observation with SKA1 MTBD in isolation **(Figure 4B)**, we found that microtubule coating was mostly uniform with no clear boundary between microtubule plus-end proximal decoration and the rest of the microtubule lattice **(Figure 4E)**. Given that FL Ska is a dimer (Huis in ’t Veld et al., 2019; Jeyaprakash et al., 2012; Welburn et al., 2009), we then dimerised the SKA1 MTBD using a short leucine zipper (LZ) motif from Gcn4, attached N-terminally to a construct containing SKA1 linker and MTBD (LZ-LMTBD, **Figure 4B, Supplementary Figure 4A)**. Based on its migration on a size-exclusion column, we conclude that LZ-LMTBD is indeed a dimer **(Supplementary Figure 4B)**. Repeating the same envelope formation assay we found that LZ-LMTBD construct forms envelopes similarly to FL Ska and Ska^SKA3Δ C^ **(Figure 4F)**. We thus conclude that the minimal Ska construct that recapitulates the interaction of a FL Ska with a microtubule is a dimer of SKA1 MTBD. Several lysine and arginine residues within the SKA1 MTBD were previously reported to reduce microtubule binding *in vitro* and extend or arrest mitotic progression *in vivo* (Abad et al., 2014; Monda et al., 2017). We introduced these mutations into our LZ LMTBD and performed envelope formation assays with each of these single or double mutants (K153A, R155A, K183/184A, K203/206A, K223/226A, R236A, and R24SA, **Figure 4G and Supplementary Figure 4A,C,D)**. We found that only the R236A mutation suppressed formation of non-uniform coating of SKA1 MTBD, while the other mutants formed envelopes, even when accompanied by an expected reduction in the overall density of microtubule-bound molecules **(Figure 4G, Supplementary Figure 4D)**. The R236A also suppressed envelope formation in full-length SKA complexes, although we did observe Ska^SKA1 R236A^ accumulation at the shortening microtubule ends **(Figure 4G, Supplementary Figure 4DE)**.

To quantify the observed effects on envelope formation, we measured the enrichment of Ska in the envelopes by calculating the ratio between the envelope fluorescence intensity, and the fluorescence intensity along the less bright region distal to the microtubule end **(Figure 4H)**. Using this method, we found that the only two constructs with a statistically significant reduction in the envelope/lattice intensity ratio were LZ-LMTBD R236A and Ska^SKA1 R236A^ **(Figure 4H)**. We thus conclude that Ska:Ska self-interactions on dynamic microtubules that lead to non-uniform coating of microtubules require dimerization of SKA1 MTBD, and can be disrupted by a point mutation R236A. We then performed a series of experiments to characterise the effects of the oligomerisation-deficient Ska carrying the R236A on its microtubule binding *in vitro* and *in vivo*.

### Oligomerisation-deficient Ska has intact microtubule binding in single-molecule conditions

We first tested the microtubule-binding properties of the mutants we used above, in conditions that would prevent them from forming cooperative Ska:Ska interactions. Using taxol stabilised microtubules, and lowering the concentration of Ska to 0.1-1 nM, we observed that FL Ska molecules binding to microtubules had a brightness corresponding to monomers and dimers **(Figure 5A, Supplementary Figure 5A)**. As expected, Ska^SKA3Δ C^ showed a similar brightness distribution, while the SKA1 MTBD only existed in the monomer form **(Supplementary Figure 5A)**.

We then measured the residence time of Ska molecules on microtubules in these conditions. FL Ska and Ska^SKA3Δ C^ interacted with microtubules with a wide distribution of residence times, sometimes exceeding 1 s, while the residence time of SKA1 MTBD never exceeded 0.4 s **(Figure 5B, C)**. The distribution of residence times of the FL Ska featured a slow and a fast component, with the fast component quite similar in its slope to the single slope observed for the SKA1 MTBD **(Figure 5C, Supplementary Figure 5)**. We interpreted this observation, together with the bimodal brightness distribution, as evidence that at sub-nanomolar concentrations the Ska dimer can partially dissociate into monomers **(Figure 5A,C, Supplementary Figure 5A)**. However, for simplicity we limited further analysis to measuring the fraction of residence times that were only observed for monomeric SKA1 MTBD and LMTBD constructs (short events, ≤ 0.3s), and the fraction of residence times that were never or very rarely observed for MTBD/LMTBD (long events, >0.3s, **Figure 5C)**. Using this approach, we found that the dimeric SKA1 LZ LMTBD construct bound to microtubules similarly to FL Ska **(Figure 5D, Supplementary Figure 5)**, which is consistent with our findings based on the envelope formation assay that SKA1 LZ-LMTBD faithfully recapitulates the microtubule interaction of FL Ska **(Figure 4)**.

**Figure 5.**
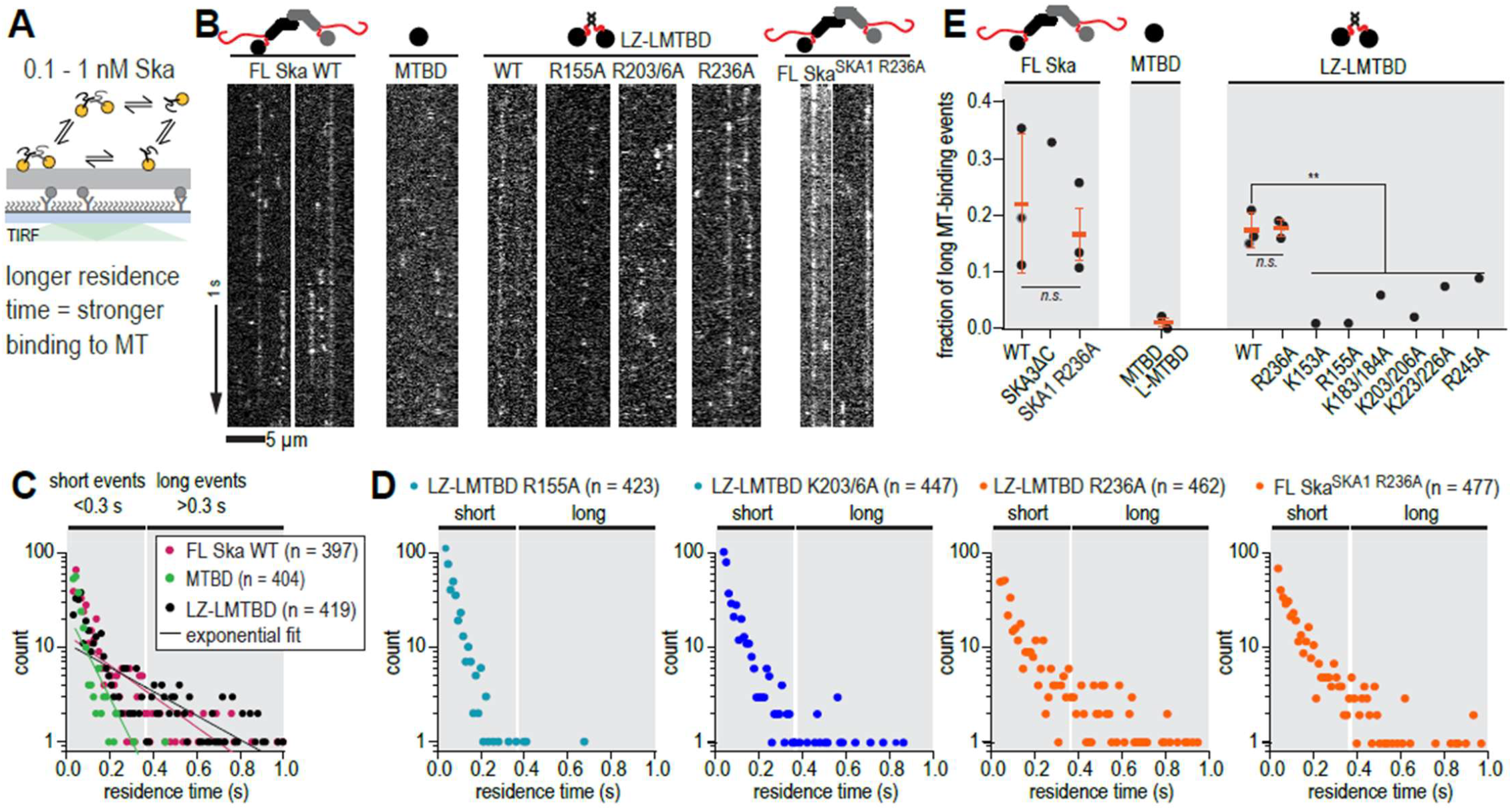
Oligomerisation-deficient Ska has intact microtubule binding in single-molecule conditions. **(A)** Schematic of the experimental setup with Ska in the concentration of 0.1-1 nM interacting with coverslip-attached taxol-stabilised microtubules. **(B)** Example kymographs showing single molecule binding events of the constructs indicated. **(C, D)** Residence time distributions of the constructs indicated. Shaded areas represent events considered ‘short’ and ‘long’ during quantification. **(E)** Fraction of ‘long’ binding events for the constructs indicated. Individual statistics per repeat are shown in Supplementary Figure 5. Orange lines show mean ± SD of repeated experiments. Welch’s t-test: FL Ska WT vs FL Ska^SKA1 R236A^ p = 0.56; LZ-LMTBD WT vs LZ-LMTBD R236A p = 0.89; FL vs LZ-LMTBD p = 0.58; FL Ska^SKA1 R236A^ vs LZ-LMTBD R236A p = 0.84; LZ-LMTBD WT vs non-R236A mutants p = 0.003. Scale bar: 5 µm (horizontal), 1 s (vertical).

We further used the LZ-LMTBD construct to screen mutations in the SKA1 MTBD for their effect on Ska residence time on stable microtubules. We observed that the majority of K-A and R-A mutations resulted in a reduction of the observed fraction of long microtubule-binding events, with the exception of LZ LMTBD R236A which bound to microtubules similarly to LZ-LMTBD **(Figure 5DE, Supplementary Figure 5)**. We confirmed that this mutation did not affect binding of single molecules of FL Ska by comparing residence times of FL Ska and FL Ska^SKA1 R236A^ **(Figure 5DE, Supplementary Figure 5)**. We thus conclude the SKA1 R236 is important for the Ska:Ska interaction, but not the Ska:microtubule interaction.

### Oligomerisation-deficient Ska fails to support cold stable microtubule attachments

Properly formed kinetochore-microtubule attachments, as opposed to most other cellular microtubules, are resistant to a brief treatment with a low temperature, an established assay used to demonstrate how kinetochores stabilise microtubule ends **(Rieder, 1981**). To assess the cold stability of kinetochore microtubules in the background of oligomerisation-deficient Ska, we depleted endogenous SKA1 using siRNA, and then expressed GFP-tagged and siRNA-resistant SKA1 in HeLa cells **(Figure 6A)**. In the absence of a commercially available anti-SKA 1 antibody, we used immunofluorescence analysis of kinetochore-localised endogenous SKA3 after treatment of cells to monitor depeletion of SKA1 by siRNA **(Supplementary Figure 6A)**. Using this method, we observed a strong reduction of kinetochore-bound SKA3 following the SKA1 siRNA treatment **(Supplementary Figure 6BC)**. Consistently, we observed an overall reduction in SKA3 levels by immunoblotting **(Supplementary Figure 6BC)**.

**Figure 6.**
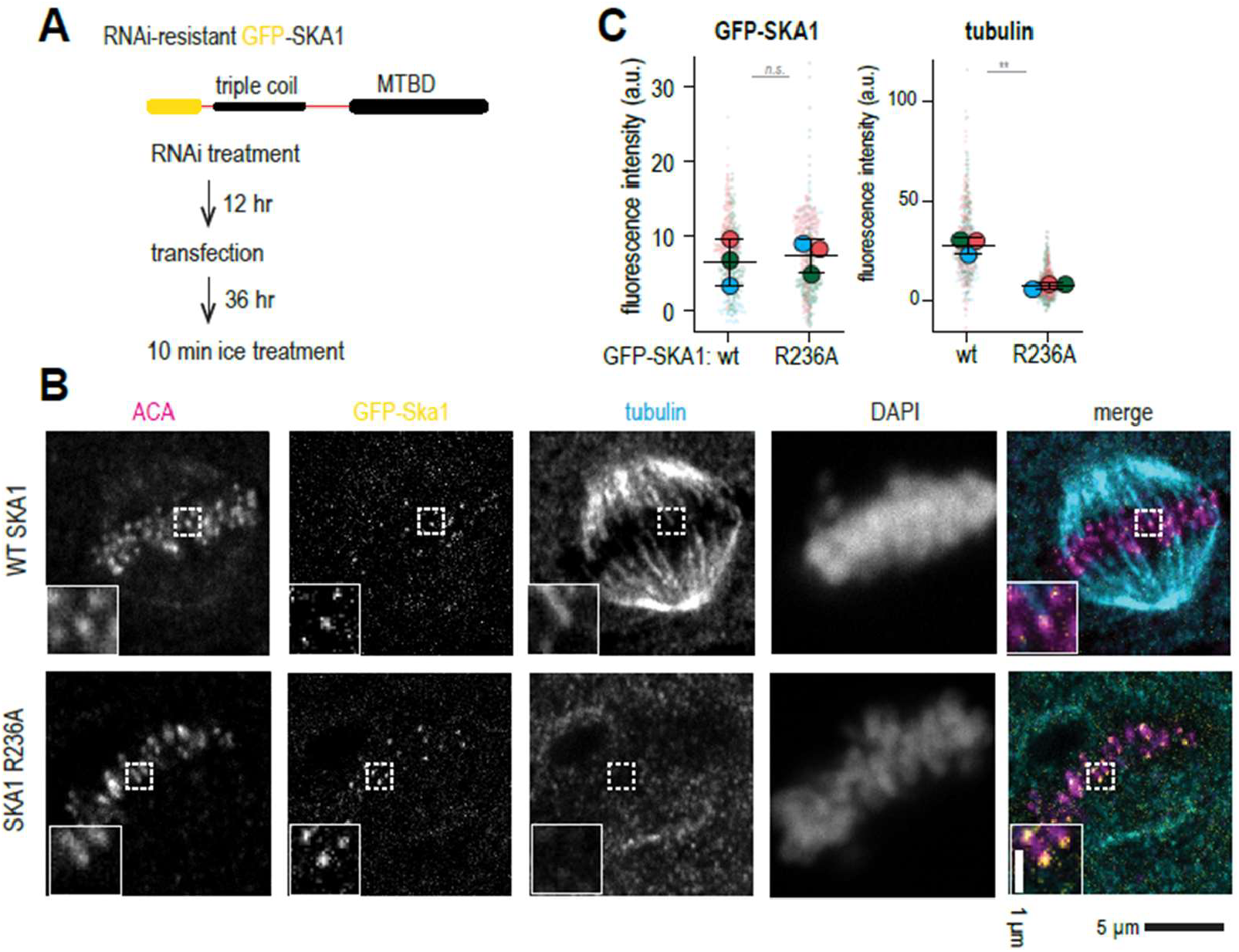
Oligomerisation deficient Ska fails to support cold-stable microtubule attachments. **(A)** Experimental setup to express RNAi-resistant GFP-SKA 1 in HeLa cells treated with RNAi to deplete endogenous SKA1, followed by the 10 minute incubation of cells on ice. **(B)** Single planes from z-stacks of confocal images of cells stained for ACA (magenta), GFP (yellow), tubulin (cyan), and DAPI. Insets show magnified regions boxed from each panel. **(C)** GFP and tubulin intensity at kinetochores. Small circles represent individual kinetochores, large circles represent mean values per repeat, black lines show mean ± SD. N = at least 20 kinetochores per cell, 22 (WT) or 23 (R236A) cells in total, repeated 3 times. Paired t-test for repeated measurements (n = 3): p = 0.0064.

Importantly, SKA3 levels were restored upon the expression of siRNA-resistant GFP-SKA 1. We have thus established a system to assess the effects of SKA1 mutants.

Incubation of cells transiently expressing WT GFP-SKA 1 on ice for 10 minutes depolymerised majority of microtubules, but kinetochore-bound microtubules were retained **(Figure 6B)**. On the contrary, expression of GFP-SKA 1 R236A followed by the same cold treatment resulted in a strong reduction, or a complete absence of kinetochore-microtubule attachments, despite GFP-SKA 1 R236A being faithfully recruited to kinetochores **(Figure 6B)**. Cells transfected with WT GFP-SKA 1 but with poor expression of the protein, as judged by the lack of the GFP signal at kinetochores, were also characterised by unstable kinetochore microtubule attachments following the cold treatment **(Supplementary Figure 6D)**. We further confirmed that despite an unchanged GFP signal at kinetochores colocalised with the ACA staining **(Figure 6C**, p = 0.77), expression of GFP-SKA 1 R236A resulted in severely reduced fluorescence intensity of kinetochore-proximal tubulin staining **(Figure 6D**, n = 3 repeats, p = 0.0064).

### Oligomerisation-deficient Ska fails to stabilise microtubule ends against disassembly *in vitro*

Finally, we used the *in vitro* microtubule-stabilisation assay we designed earlier **(Figure 7A)** to test the effect of SKA 1 R236A mutation. We repeated the experiment with FL Ska WT and FL Ska^SKA1 R236A^, and found that 1 nM FL Ska^SKA1 R236^, in presence of 1 nM Ndc80, failed to stabilise microtubules against disassembly, unlike FL Ska WT in similar conditions **(Figure 7B)**. Notably, the oligomerisation-deficient FL Ska^SKA1 R236^ was able to follow shortening microtubule ends in this experiment, as we also observed earlier in conditions that did not force microtubules to depolymerise **(Figure 4G, 7B)**. As a consequence of the oligomerisation-deficient mutant’s inability to stabilise microtubule ends, we did not observe an enrichment of fluorescent microtubule extensions following 10 minutes oftubulin dilution in the presence of Ndc80 and FL Ska^SKA1 R236A^, compared to Ndc80 alone (2-way ANOVA: p = 0.11; **Figure 6C)**. We thus conclude that Ska oligomerisation promotes cooperative Ska:Ndc80 interactions, thereby preventing disassembly of kinetochore-attached microtubules of the mitotic spindle.

**Figure 7.**
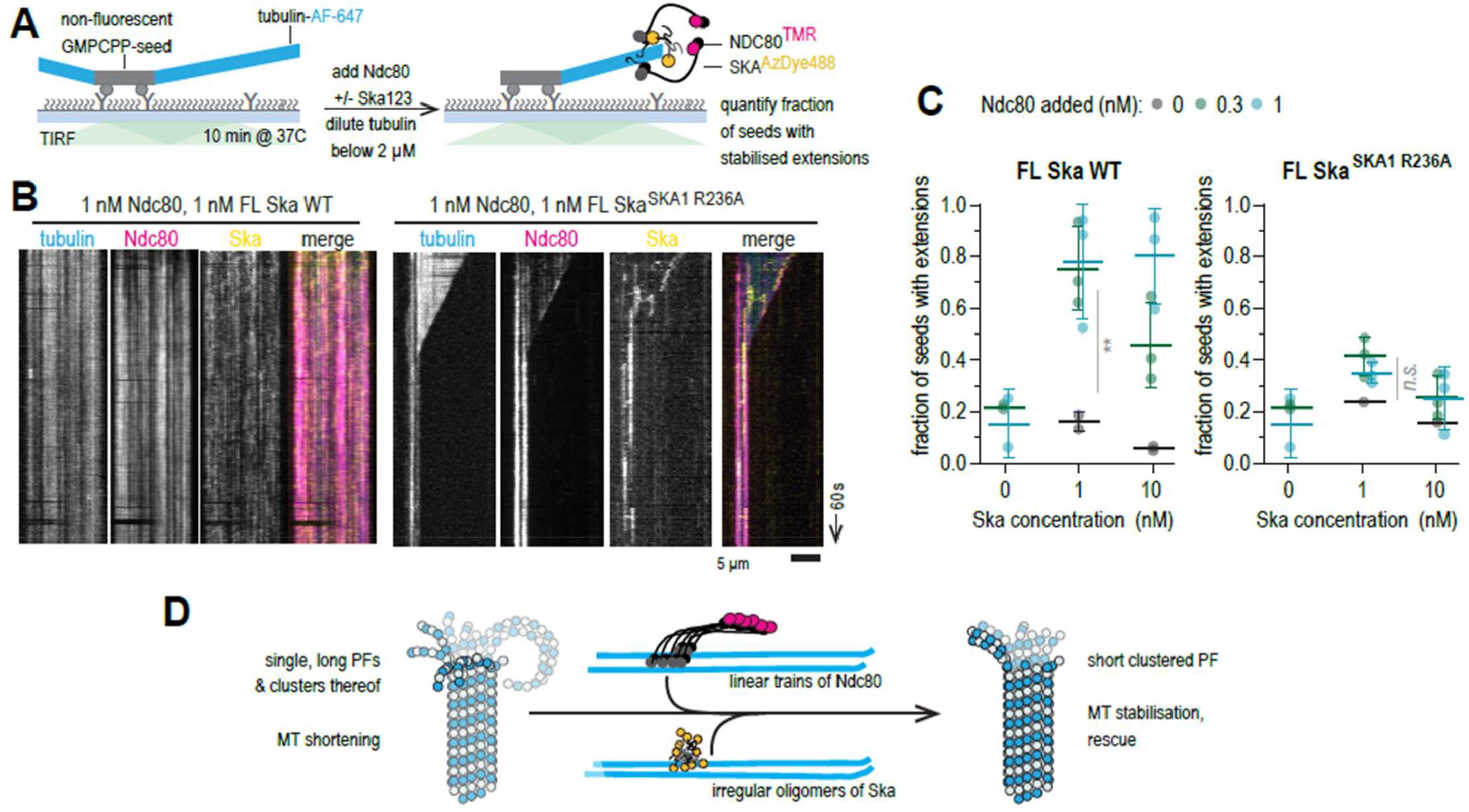
Oligomerisation-deficient Ska fails to stabilise microtubule ends against disassembly *in vitro*. **(A)** Schematic of an experiment to test microtubule end stabilisation by Ndc80 and Ska. **(B)** Example kymographs showing microtubule length over time following the simultaneous addition of 1 nM Ndc80 with 1 nN Ska (WT or SKA1 R236A). **(C)** Fraction of unlabelled GMPCPP stabilised microtubule seeds with fluorescenttubulin extension following 10 minutes aftertubulin dilution. Grey: no Ndc80 added, green: 0.3 nM Ndc80 added, cyan: 1 nM Ndc80 added (at least 50 seeds quantified in total over at least 5 fields of view per repeat, repeated 2-3 times). Lines and error bars represent mean± SD. Two-way ANOVA for FL Ska WT: row factor (Ska concentration) p = 0.0012; column factor (Ndc80 concentration) p = 0.0002. Two-way ANOVA for FL Ska^SKA1 R236A^ row factor (Ska concentration) p = 0.0084; column factor (Ndc80 concentration) p = 0.11. **(D)** Cooperative oligomers of Ndc80 and Ska promote microtubule stabilisation via clustering oftubulin protofilaments.

## DISCUSSION

The binding between outer kinetochores and microtubule ends is highly multivalent and confined in space and time to the middle of a mitotic cell. It is therefore tremendously difficult to systematically disentangle the interactions of microtubule-binders with microtubules from hetero-or homotypic interactions between microtubule-binders. Here, we developed an experimental strategy to distinguish Ska:Ska interactions on microtubules from direct Ska:microtubule binding, by enabling rapid screening of point mutants of SKA1 MTBD using a minimal construct that recapitulates the FL Ska microtubule-binding properties. Comparing residence time of SKA1 MTBD mutants in single molecule conditions to uniformity of their microtubule decoration at high concentrations, we have identified the R236A mutation as the one that impairs the normally observed non-uniform microtubule decoration at high Ska concentrations without affecting individual molecules’ residence time on microtubules **(Figures 4-5)**. We further confirmed that this mutation has similar effects on Ska:Ska and Ska:microtubule interactions both in the minimal LZ-LMTBD construct, and in the FL Ska complex.

A number of alanine substitutions of lysine and arginine residues, in various combinations, were reported to affect the amount of Ska sedimented after centrifugation with microtubules, the microtubule end tracking properties of purified Ska, as well as the duration of mitotic progression **(Abad et al., 2014; Monda et al., 2017)**. This also applies to the R236A mutation, which was shown to reduce bulk microtubule binding and extend the duration of mitosis, alone and in combination with R24SA and R1 SSA **(Abad et al., 2014; Monda et al., 2017)**, although without major mitotic alignment defects. Results presented here allow us to conclude that contrary to the rest of the R/A and K/A mutations tested, which affect the Ska:microtubule interaction directly, the R236A mutations exerts its phenotype through impaired Ska:Ska interactions. It is important to note that FL Ska^SKA1 R236A^ in our *in vitro* experiments was able to follow the shortening microtubule ends **(Figure 4G, 7B)**, further reinforcing our interpretation that this mutation does not affect Ska’s ability to interact with microtubule lattices or ends per se.

We further showed that SKA1 R236A, despite interacting with microtubules normally in vitro, led to a severe reduction in the cold-stability of kinetochore fibers, a hallmark of reduced kinetochore-microtubule binding strength **(Figure 6)**. This experiment is directly supplemented by our *in vitro* test for microtubule end stabilization by the cooperative oligomers of Ndc80 and Ska **(Figure 7)**. Ndc80 and Ska at low nanomolar concentrations stabilize microtubules against disassembly triggered by sudden dilution of tubulin, however substitution of WT Ska to Ska^SKA1 R236A^ leads to loss of this stabilisation.

There is no high-resolution structure of microtubule-bound Ska available. Results of our analysis of individual microtubule-bound oligomers of Ska in tomograms **(Figure 3F)** are consistent with the formation of multi-layered Ska oligomers, that are held together, at least partially, by Ska:Ska interactions, because their dimensions exceed the expected dimensions of a single layer of microtubule-bound Ska. These multi-layered assemblies are not typical for microtubule-bound proteins: we only observed a single microtubule-bound layer of Ndc80 oligomers **(Figure 2)**, and the same was true for a previously described microtubule plus-end tracking protein network, which formed comets that were also consistent with a single layer microtubule decoration **(Maan et al., 2023)**.

Given that Ska was previously reported to interact with soluble tubulin (Monda et al., 2017), we can not rule out whether SKA1 R236 is involved in a direct interaction with another residue of SKA1 MTBD, or is part of an indirect binding of some kind. Our efforts to use subtomogram averaging to gain a higher resolution structure only allowed us to conclude that, under the conditions tested, Ska does not interact with microtubules in a regular, ordered fashion **(Figure 3)**. It is possible that Ska:Ska self-interactions stabilise microtubule ends by cross-linking neighbouring tubulin protofilaments and prevents them from splaying apart. Alternatively, if SKA1 MTBDs oligomerise along the length of the same protofilament, like the Ndc80 complex **(Figure 2-3)** (Alushin et al., 2010), they might affect the protofilament curvature. Our recent comparison of the structures of growing and shortening microtubule ends predicts that both lateral protofilament cross-linking, as well as individual protofilament straightening have a similar effect on converting a shortening microtubule end into a configuration compatible with growth **(Kalutskii et al., 2025)**. Direct comparison of microtubule plus-ends stabilised against shortening by Ndc80 alone, or by Ndc80:Ska oligomers, demonstrates that both treatments promote protofilament clustering **(Figure 2)**. While it is difficult to disengage effects of protofilament straightening and lateral clustering, the similarity of the observed changes in the shapes of microtubule plus ends suggests that the main stabilising effect is exerted through the oligomerisation of Ndc80. This conclusion is corroborated by our low-resolution averages of “trains” of Ndc80 CH-domains on two neighbouring protofilaments, in all conditions tested, with and without Ska present **(Figure 3AB)**. In this context, we propose that Ska may be required to further oligomerise Ndc80 when its concentration or availability is insufficient, or else to shift Ndc80 “trains” from the microtubule lattice towards microtubule ends. We thus conclude that a combination of Ndc80 oligomerisation via the loop region and the microtubule-bound CH-domains (Alushin et al., 201O; Polley et al., 2023), and Ska cross linking via the SKA1 MTBD, and potentially also SKA3 tail, are all necessary to achieve the required level of end stabilisation **(Figure 7D)**.

The kinetochore-microtubule binding has been previously shown to be stabilised by the applied force (Akiyoshi et al., 2010; Nicklas and Ward, 1994). We previously demonstrated how presence of Ska at the interface between the Ndc80 oligomer and a shortening microtubule end during force production can stabilise the stalled conformation of the microtubule ends, and in this way to increase the rate of force-dependent microtubule rescue **(Huis in ’t Veld et al., 2019)**. Our direct imaging of microtubule ends stabilised by Ndc80:Ska oligomers brings new information about the functioning of this system, such as the lateral clustering or protofilaments by Ndc80, and the effect of Ska on the shortening ofthe protofilaments **(Figure 2)**. Future work will be necessary to understand how the structures of microtubule ends, and Ndc80 and Ska oligomers binding to them, are affected by the applied force.

It also remains to be tested whether Ska recruitment to the interface between the Ndc80 and the microtubule end is force sensitive. Although our previous results with isolated Ndc80 trimers argue against this possibility **(Nick Maleki et al., 2023)**, higher order oligomers of Ndc80, such as observed in this study, can recruit Ska more efficiently in response to the tension generated across the kinetochore-microtubule interface. Thus, it still remains unclear what triggers an increased recruitment of Ska to the kinetochore so late before the metaphase-to-anaphase transition **(Auckland et al., 2017; Cheerambathur et al., 2017)**, since the only other known determinant of the direct Ska:Ndc80 interaction, namely the Cdk1 phosphorylation of SKA3 T358 and T360 **(Huis in ’t Veld et al., 2019; Zhang et al., 2017)**, should be present throughout mitosis due to high levels of Cdk1 activity. It remains to be tested whether effects of MPS1 **(Maciejowski et al., 2017)**, PP1 **(Conti et al., 2019; Sivakumar et al., 2016)**, or other regulatory events affect the assembly of cooperative Ndc80:Ska oligomers at the correct time after the kinetochore biorientation.

## ACKNOWLEDGMENTS

VAV is grateful to Andrea Musacchio (Max-Planck Institute of Molecular Physiology) and Marileen Dogterom (Delft University of Technology) for supporting the initial phase of this work. We also thank Charlotte Millership (insect cell facility, OMUL) for help with insect cell expression, Hui Zhang (EM facility, OMUL) for help with grid preparation and screening, Christoph Diebolder (NeCEN}, Nora Cronin (LonCEM) and Emma Buzzard (Diamond Light Source) for help with cryoET data collection, Michaela Egertova (OMUL) for help with confocal microscopy. We thank Antony Oliver (University of Sussex) for a gift of a plasmid encoding for Cdk1:CyclinB:CKS1, Viji Draviam (OMUL) for a gift of a plasmid encoding for GFP-PMF1, and Michel Steinmetz (Paul Scherer Institute) for a gift of a plasmid encoding for MACF-GCN4. VAV acknowledges grant support from the Biotechnology and Biological Sciences Research Council (BB/X014975/1) and the Wellcome Trust (308895/Z/23/Z). Confocal imaging using the Leica Stellaris 8 system was supported by BBSRC grant BB/W019698/1.

## SUPPLEMENTARY MATERIALS

Supplementary Figures 1-6.

## MATERIALS AND METHODS

### Cloning, expression, and purification of full length Ndc80 and Ska complexes

Full-length Ndc80 and Ska complexes were expressed using previously described vectors (Huis in ’t Veld et al., 2019; Volkov et al., 2018). R236A mutation was introduced into a pLIB vector containing SKA1 using conventional site directed mutagenesis techniques and a pBIG1 vector containing SKA3, SKA2, and SKA1 R236A was generated using Gibson assembly. Baculovirus generation, and protein expression were both carried out in *Sf9* insect cells. Cells were harvested 2-3 days after infection, washed in PBS and stored at −80C. Cell were thawed, resuspended in lysis buffer (20 mM Tris-HCI, pH 8.0, 150 mM NaCl, 5% v/v glycerol, 2 mM DTT, 20 mM imidazole, 0.5 mM PMSF, and protease-inhibitor mix (Pierce), and DNAse I (Roche)), lysed in a dounce glass homogenizer on ice (using 15-30 passes of the pestle), and the lysate was cleared by centrifugation at ca. 88,000 g for 45 min. The cleared lysate was mixed with pre-equilibrated HIS Select Ni affinity beads (Merck) and incubated for 1-2 hrs at 4C with rotation. Beads were washed using 50 ml of lysis buffer without protease inhibitors, collected in a gravity flow column, and the protein was eluted manually in 1 ml fractions using the same buffer with 250 mM imidazole.

FL Ska WT was further purified by ion-exchange chromatography. Ni-purified fractions were diluted 5-fold in buffer A (20 mM Tris-HCI, pH 8.0, 30 mM NaCl, 5% v/v glycerol, 1 mM DTT), applied to a Capto HiRes0 5/50 column (Cytiva) equilibrated in the same buffer, and eluted using a linear gradient from 30 to 500 mM NaCl in 80 ml. This step was omitted for FL Ska^SKA1 R236A^ which eluted from Ni beads in a clean enough form, and the protein was instead desalted using a PD-10 desalting column. In both cases, relevant fractions were pooled, concentrated in 50 kDa molecular weight cut-off concentrators (Thermo Fisher) and applied to a Superose 6 Increase 10/300 column (Cytiva) equilibrated in 20 mM Tris-HCI, pH 8.0, 150 mM NaCl, 5% v/v glycerol, 1 mM DTT. Relevant fractions were pooled, concentrated, flash-frozen in liquid nitrogen, and stored at −80°C.

FL Ndc80 was expressed and purified as described previously (Huis in ’t Veld et al., 2019) with slight modifications. Baculovirus generation and protein expression were both carried out in *Sf9* insect cells. Cells were resuspended in lysis buffer (50 mM Hepes, pH 8.0, 250 mM NaCl, 5% v/v glycerol, 1 mM DTT, 20 mM imidazole, 0.5 mM PMSF, and protease-inhibitor mix (Pierce)), lysed in a dounce glass homogenizer on ice (using 15-30 passes of the pestle), and the lysate was cleared by centrifugation at ca. 88,000 g for 45 min. Ni affinity and SEC were performed as described above using a buffer with 50 mM Hepes and pH 8.0, 250 mM NaCl 5% v/v glycerol, 1 mM DTT. For separation on a Capto HiRes O 5/50 column, protein was eluted with a linear gradient from 25 to 300 mM NaCl in 40 ml.

To label the proteins fluorescently, the C terminal 6xHis tag on SKA1 was replaced with a fluorescent peptide GGGK-AzDye488 (Vector labs), or the C-terminal 6xHis tag on SPC24 was replaced with a fluorescent peptide GGGGK-TMR (Thermo Fisher) in a 30 min reaction at 25C, immediately followed by size exclusion chromatography. Sortase 7M (Hirakawa et al., 2015), the protein complex, and the peptide were used in an approximate molar ratio of 1:10:100.

### Purification of Cdk1 and in vitro phosphorylation of Ska

Cdk1, Cyclin B, and CKS1 were coexpressed from the same vector (a kind gift from Antony Oliver) in *SF9* insect cells. Cell pellets were resuspended in the lysis buffer containing 25 mM HEPES pH 7.5, 200 mM NaCl, 0.5 mM TCEP, protease inhibitor cocktail and DNAse, lysed by sonication, and the lysate was cleared by centrifugation at 36,000 g for 60 minutes. Cleared lysate was applied to a 5ml HiTrap TALON FF column, washed with buffer A containing 25 mM HEPES pH 7.5, 200 mM NaCl, 0.5 mM TCEP, then the same buffer containing 5 mM imidazole, and finally eluted in 25 ml of the same buffer containing 250 mM imidazole in a single step. Eluted material was diluted two-fold with lysis buffer and applied to a 1ml HiTrap StrepXT column. The column was washed with buffer A, and the protein was eluted in 5 ml of the same buffer containing 50 mM Biotin in a single step. Strep elution fraction was concentrated to 500 ul and loaded onto a Superdex 200 10/300 column equilibrated in 10 mM HEPES pH 7.5, 200 mM NaCl, 0.5 mM TCEP, 5% (v/v) glycerol, 0.002% Tween-20. Relevant fractions were pooled, flash-frozen in liquid nitrogen and stored at *-BOC*.

For *in vitro* phosphorylation, Ska complexes were exposed to CDK1:Cyclin-B in presence of 5 mM ATP and 10 mM MgCb for 30 min at 25C, followed another 30 min with the addition of Sortase and fluorescent peptide GGGK-AzDye488 in an approximate ration of 1/10 and 1Ox to the Ska, respectively. Phosphorylated and labelled Ska was immediately separated from the rest of components using Superose 6 Increase 10/300 column (Cytiva) equilibrated in 20 mM Tris-HCI, pH 8.0, 150 mM NaCl, 5% v/v glycerol, 1 mM DTT. Relevant fractions were pooled, concentrated, flash-frozen in liquid nitrogen, and stored at −80°C.

### Cloning, expression, and purification of SKA1 fragments

Fragments of SKA1 were subcloned from pLIB-SKA 1 into pET28a+, and the Gcn4 leucine zipper fragment RMKOLEDKVEELLSKNYHLENEVARLKKLVG ER was inserted using two rounds of PCR. Site directed mutagenesis of SKA1 MTBD was performed in pET28a+ using conventional methods.

Expression of SKA 1 fragments was performed in *e*.*coli* BL21(DE3) Rosetta cells grown at 37C in presence of Chloramphenicol and Kanamycin to an OD600 of 0.6. Protein expression was induced by the addition of 200 mM IPTG, and continued for 14-20 hrs at 18C. Cells were washed in PBS and pellets were stored at *-BOC*. Cells were thawed on ice and resuspended in a lysis buffer (20 mM Tris-HCI, pH 8.0, 150 mM NaCl, 5% v/v glycerol, 1 mM DTT, 20 mM imidazole, 0.5 mM PMSF, and protease-inhibitor mix (Pierce}, and DNAse I}, lysed by sonication and cleared by centrifugation at 75,000 g. The cleared lysate was mixed with pre-equilibrated HIS-

Select Ni affinity beads (Merck) and incubated for 1-2 hrs at 4C with rotation. Beads were washed three times using lysis buffer without protease inhibitors, collected in a gravity flow column, and the protein was eluted manually in 1 ml fractions using the same buffer with 400 mM imidazole. Peak fractions were desalted using a PD-10 desalting column and concentrated in 10 kDa molecular weight cut-off concentrators (Thermo Fisher).

To label the proteins fluorescently the C terminal 6xHis tag on SKA1 was replaced with a fluorescent peptide GGGK-AzDye488 (Vector labs) using sortase in a 30 min reaction at 25C as described above, immediately followed by size-exclusion chromatography using a Superdex 75 Increase 10/300 column pre-equilibrated in 20 mM Tris-HCI, pH 8.0, 150 mM NaCl, 5% v/v glycerol, 1 mM DTT. Relevant fractions were pooled, concentrated in 10 kDa molecular weight cut-off concentrators, aliquoted and stored at −80C.

### Tubulin, microtubules, and preparation of flow chambers

Porcine brain tubulin was purified and labelled in house using standard protocols (Castaldi and Popov, 2003; Hyman et al., 1991) using amine reactive Digoxigenin (Merck) or AlexaFluor-647 (lnvitrogen). DIG-labelled GMPCPP-stabilised seeds were produced using two cycles of polymerization to fully replace GTP, as described previously (Volkov et al., 2018).

Taxol-stabilised microtubules were made by incubating 70 **µM** tubulin (with or without DIG- and fluorescent labels) in presence of 25% glycerol and 1 mM GTP for 20 min at 37°C, and then stabilized by an addition of 25 **µM** taxol followed by another 20 min incubation. Polymerised microtubules were sedimented by centrifugation at 100,000 g in a TLA100 rotor over 60% glycerol cushion, and the pellet was resuspended in a buffer containing 80 mM K-PIPES pH 6.9, 1 mM EGTA, 4 mM MgCb, supplemented with 40 µM taxol. Polymerised microtubules were stored at 25°C for up to 3 days.

Glass coverslips were silanised using PlusOne Repel Silane (Cytiva}, coated with anti-DIG antibody (Roche), and passivated using 1% Pluronic F-127 (Merck) as described previously (Volkov et al., 2018). All experiments with dynamic microtubules were performed using the following buffer: 80 mM K-PIPES pH 6.9, 1 mM EGTA, 4 mM MgCl_2_, 1 mM GTP, 1 mg/ml κ-casein, 0.1% methylcellulose, oxygen-scavenging mix consisting of 4 mM DTT, 0.2 mg/ml catalase, 0.4 mg/ml glucose oxidase and 20 mM glucose, 10-11 µM tubulin (3-5% fluorescently labelled), at 30C. All experiments with taxol-stabilised microtubules were performed using the following buffer: 80 mM K-PIPES pH 6.9, 1 mM EGTA, 4 mM MgCb, 1 mg/ml κ-casein, 0.1% methylcellulose, 40 µM taxol, oxygen-scavenging mix consisting of 4 mM DTT, 0.2 mg/ml catalase, 0.4 mg/ml glucose oxidase and 20 mM glucose, at 25C.

Microtubule end-stabilisation experiments were performed by first growing microtubules in a flow chamber using a solution containing 80 mM K-PIPES pH 6.9, 1 mM EGTA, 4 mM MgCl2, 1 mM GTP, 1 mg/ml κ-casein, 0.1% methylcellulose, oxygen-scavenging mix consisting of 4 mM DTT, 0.2 mg/ml catalase, 0.4 mg/ml glucose oxidase and 20 mM glucose, 10 µM tubulin (3% fluorescently labelled), at 37C. After 10 min of incubation, the solution was exchanged to the one containing the same buffer without soluble tubulin (pre warmed at 37C), but with addition of Ndc80 and/or Ska in the required concentration, the chamber was then sealed with VALAP, placed on the microscope stage pre warmed to 30C, and imaging was commenced immediately. The time between the change of solution and start of imaging was typically 30-60 s.

### TIRF microscopy

Experiments were performed using a custom microscope based on the OpenFrame system (Lightley et al., 2023) and manufactured by Cairn Research. The instrument is equipped with the following lasers: a 150 mW Omicron LuxX 488 nm, a 200 mW Omicron LuxX 638 nm, and a 150 mW OBIS 561 nm, steered through the GATACA llas^2^ TIRF/FRAP illumination module into a Nikon CFI Apochromat TIRF 100x objective with NA1.49. The other components include an ASI motorized XY stage, a Cairn MonoLED for brightfield imaging, a custom Cairn autofocus system, and an Okolabs temperature controller with an objective heating collar. Images were acquired using an ET 405/488/561 /640nm Laser Quad Band Set (Chroma) and an iXon Life 897 EMCCD (Andor) using the MicroManager 2.0 software (Edelstein et al., 2014).

### Electron cryo-tomography

Samples with dynamic microtubules decorated with Ska were prepared as described previously (Maan et al., 2023). SiO holey grids (SPI supplies) were treated with oxygen plasma and immediately immersed in PlusOne Repel Silane solution (Cytiva) for 2 min, followed by sequential rinsing in Ethanol and water. Each grid was taken by Leica EM GP2 plunger forceps, incubated for 1 min with a 7 uL drop of 0.2 µM anti-DIG antibody (Roche), and sequentially rinsed with MRB80, 1% Pluronic F-127, and MRB80. A grid passivated in this way was put in the chamber of the plunger equilibrated at 25C and 95% humidity, and further incubated with 9 µL of GMPCPP stabilised microtubule seeds, followed by rinsing and addition of 5 µL of a buffer containing 25 **µM** tubulin, **1** mM GTP, and 1 mM DTT in 80 mM K-PIPES pH 6.9, 1 mM EGTA, 4 mM MgCb. Microtubules were allowed to grow for 5 min, after which the solution on the grid was replaced with the one containing 400 nM FL Ska and 5 nm gold particles, in addition to 25 µM tubulin, 1 mM GTP, and 1 mM DTT. Grids were blotted from the back side, plunge-frozen in liquid ethane, and stored in liquid nitrogen.

Tilt series were recorded using a Titan Krios microscope (FEI) equipped with a Gatan K2 electron detector (NeCEN). Automated image acquisition was performed using Tomography software (Thermo Fisher). Images were recorded at 300 kV with a nominal magnification of 33,000x, resulting in the pixel size of 4.24 Å at the specimen level. Energy filtering was performed at post-processing using a 30 eV-wide slit. Bi directional tilt series ranged from 0° to ±60° with a tilt increment of 2°, with the total electron dose of 100 e^−^/Å^2^ andthetarget defocus set to 4 µm.

Samples with Ndc80 and Ska oligomers stabilising the shortening microtubules were prepared using SiO Ouantifoil R2/2 grids (EMS) passivated as described above. Microtubules were grown on grids suspended in the chamber of a Leica EM GP2 plunger equilibrated at 37C and 95% humidity. Microtubules were grown for 10 min in presence of the buffer containing 80 mM K-PIPES pH 6.9, 1 mM EGTA, 4 μM

MgCl_2_, 12 µM tubulin, 1 mM GTP, 1 mM DTT, and 1 mg/ml κ-casein. After this incubation, the ca. 4 µL of solution on the grid was mixed by pipetting with ca. 30 µL of a pre-warmed solution containing the same components, with tubulin substituted to Ndc80 +/-Ska at a given concentration, and 5 nm gold nanoparticles. Following the mixing, majority of the solution was removed, leaving 3-4 µL on the grid. After a 2 min incubation in the chamber of the Leica EM GP2 plunger, the grids were blotted from the back side and plunge frozen in liquid ethane. Tilt series were recorded using Titan Krios microscopes (FEI) equipped with a Gatan K3 electron detector. Two grids with 3 and 10nM Ndc80 respectively were imaged at LonCEM (London Consortium EM, Francis Crick Institute), with data acquisition using TOMO5 software (FEI), at 300 kV with 42,000x magnification resulting in the pixel size of 2.11 A at the specimen level. One grid with Ndc80 and Ska was imaged at the National Cryo-EM facility at eBIC, located within the Diamond Light Source, using TOMO5 software (FEI), at 300 kV with 53,000x magnification resulting in the pixel size of 1.63 Å at the specimen level. In all cases, bi-directional tilt series were recorded using a dose-symmetric scheme in the range from 0° to ±60° with a tilt increment of 3°, the total electron dose of 100 e-/Å^2^ andthe target defocus set in a range from 2.5 to 4.5 µm.

Tomograms were reconstructed, denoised, and analysed for microtubule polarity and oligomer dimensions as described previously **(Maan et al., 2023)**, with motion correction using MotionCor2 **(Zheng et al., 2017)**, tilt series alignment and back projection of tomograms using IMOD **(Kremer et al., 1996)**, and denoising using cryoCARE **(Buchholz et al., 2019)**. Protofilament length and clustering were determined as described previously (Kalutskii et al., 2025).

### Subtomogram averaging

Unprocessed raw movies were re-processed using the tomography workflow in Relion 5.0 (Burt et al., 2024). Motion correction was performed using MotionCor2, CTF correction was performed using CTFfind 4.1 (Rohou and Grigorieff, 2015) and tomograms were reconstructed in an automated mode using IMOD. Particles were picked as filaments with a repeat of 4.1Å, using denoised tomograms in napari (Sofroniew et al., 2025). Microtubule polarity during particle picking was determined using inflection of the Ndc80 coiled coils, which pointed towards microtubule plus-ends. Subtomograms were extracted using a binning factor 4, and a box size of 192 unbinned pixels, then cropped to 96 unbinned pixels. Initial models were generated de novo using the variable-metric gradient descent algorithm over 100 iterations, without masking, and without any prior orientation of particles, using **C1** symmetry. Initial models generated this way were used for automated 3D refinement, which was followed by removal of duplicate particles, and 3D classification and additional rounds of 3D refinement. In all samples, with or without addition of Ska, we did not find separate classes of microtubule-decorating proteins within a sample, over several rounds of classification with various number of classes from 3 to 9, and with various masks intended to dampen the contribution of the microtubule signal. Particles obtained from Ndc80-only samples were further refined with binning 2. Particles obtained from the sample with both Ndc80 and Ska present were only refined at binning 4. 3D rendering of post processed particle averages was performed in Chimera X 1.8 (Pettersen et al., 2021).

### In vivo expression of SKA1

Hela cells were grown in DMEM containing 4.Sg/L Glucose, 2mM L-Glutamine, 110 mg/L sodium Pyruvate supplemented with 10% FBS and 100 µg/ml each of penicillin and streptomycin. Cultures were maintained at standard conditions at 37 °C in a humidified incubator with 5% CO2. For SKA1 knockdown, Hela cells at approximately 65% confluence were transfected with siGENOME Ska1-targeting siRNA (sequence: 5′-CCCGCTTAACCTATAATCAAA-3′; Horizon Cat # D-015917-04). A non-targeting control siRNA (sequence: 5′-GCCAUUCUAUCCUCUAGAGGAUG-3′; Horizon Cat # D-001210-01) was used for comparison. Transfections were performed using Lipofectamine RNAiMAX (lnvitrogen).

siRNA resistant SKA1 DNA fragment was produced by gene synthesis (Genewiz). The synthesized WT SKA1 DNA fragment was further inserted into a pGFP-N-GW vector backbone amplified from a vector encoding GFP-PMF1 (kind gift from Viji Draviam). GFP Ska1 R236A mutant was generated by using site directed mutagenesis by using siRNA resistant GFP SKA1 WT as template. For rescue assays, siRNA-resistant GFP-SKA1 constructs (wild-type and the R236A mutant) were transfected into cells after 12 hours of treatment with siRNA. Plasmids were expressed for 36 hours and transfection was carried out using Lipofectamine 3000 (lnvitrogen). Microtubule cold stability assay was performed by incubating cells expressing GFP SKA1 WT and GFP SKA1 R236A mutant on ice for 10 minutes, followed by rinsing with ice-cold PBS; the cells were then immediately fixed in −20°C methanol.

### lmmunofluorescence and confocal microscopy

Coverslips fixed in −20 °C methanol were blocked and permeabilised using PBS containing 1% bovine serum albumin and 0.5% Triton X-100 for 1 hour. Primary antibody, in a dilution specified below, was incubated for 2 hours at room temperature. Fluorophore conjugated secondary antibody diluted 1:1000 were incubated for 1 hour. The coverslips were stained with DAPI for 45 seconds and mounted on glass slides using Prolong Diamond antifade reagent (lnvitrogen). The following antibodies and dilutions were used: β-Tubulin 1:1000 (Abeam #ab6046), ACA 1:1000 (Antibodies Inc. #15-234), SKA3 1:700 (Santa Cruz Biotechnology #SC 390966), anti-rabbit AlexaFluor 568 (Abeam #ab175471), anti-Human AlexaFluor-647 (Jackson #609-604-213), anti-mouse AlexaFluorr 488 (lnvitrogen #A10680).

Confocal microscopy images were obtained with a Leica Stellaris SP8 laser scanning confocal microscope using an HC PL APO CS2 63X/1.40 Oil immersion objective. Excitation was provided by a white light laser in the range of 440-790nm for GFP, AlexaFluor-488, and AlexaFluor-568, and a diode laser 405 nm for DAPI. Fluorescence emission was collected from 430-484 nm for DAPI; 494-573 nm for GFP and AlexaFluor-488; 585-643 nm for AlexaFluor-568 and 663-800 nm for AlexaFluor-647. The confocal pinhole was set as airy 1 for all images. Optical sections in the z direction of ∽ 0.25 µm, and images were acquired at a scan speed of 400 Hz with a pixel dwell time of 2.825 µs, 2 × line averaging and a 512 × 512 pixel format.

### Image analysis

All image analyses were performed in Fiji **(Schindelin et al., 2012)**. Kymographs were generated using a custom lmageJ script (https://github.com/volkovdelft/kymo.jl). Lifetimes in single-molecule conditions were measured manually in kymographs. Ratios of envelope intens1t1es were measured using boxed regions in kymographs and calculated as *I*_*envelope*_ − *I*_*BG*_*) /(I*_*Jattice*_ − *I*_*BG*_), where *I*_*BG*_ represents background intensity next to the microtubule, measured using the same region of interest (ROI).

Intensities of SKA **1**, SKA3, and tubulin at kinetochores were measured in individual Z-planes of confocal imaging stacks, by manually selecting ROls encompassing the ACA signal and projecting several pixels towards the relevant spindle pole, to capture the end of a k-fiber. The same ROI was used for SKA and tubulin channels. At least 20 kinetochores were quantified per cell and averaged to create a mean value per cell, which were in turn averaged to generate a mean value per repeat. Statistical analyses were performed in GraphPad Prism 10.

### Statement on the use of Al

The use of machine learning was limited to cryoCARE denoising of tomograms. No other Al or ML algorithms were used for image generation or analysis. At no point in this study artificial intelligence or large language models were used for literature review, logical organization, writing, data analysis, and proofreading.

