## Supplementary Figures 1-6 for "Microtubule end stabilisation by cooperative oligomers of Ska and Ndc80 complexes"

### Supplementary Figure 1.

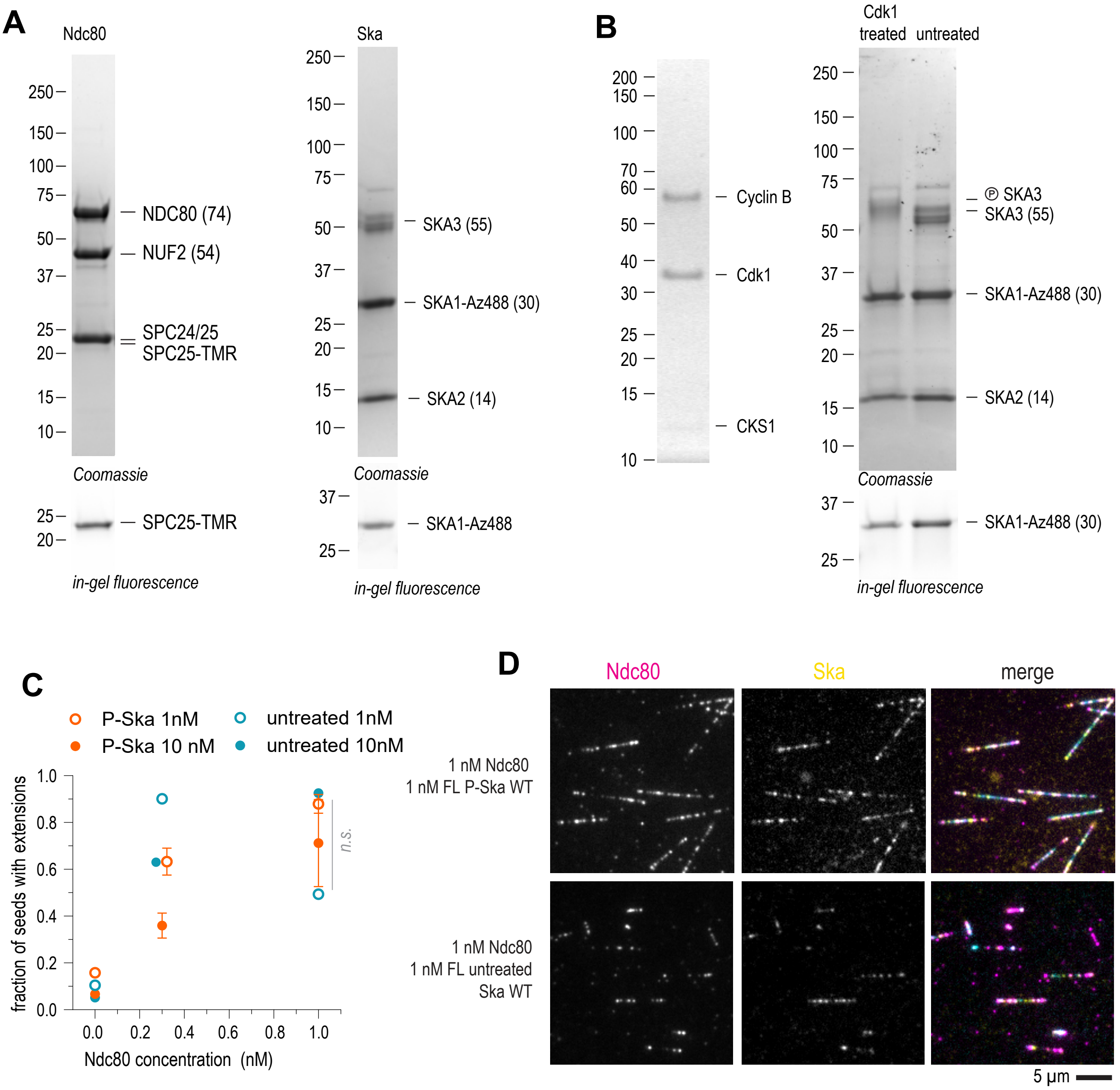


(A) SDS-PAGE of full-length Ndc80 and Ska complexes, using Coomassie staining (top), or in-gel fluorescence of TMR (left) or AzDye-488 (right). (B) SDS-PAGE of the CyclinB/Cdk1/CKS1 complex (left) and Ska complex treated with CyclinB/Cdk1/CKS1 next to an untreated control. (C) Comparison of the microtubule-stabilising activity of Cdk1-treated and untreated Ska in presence of an indicated concentration of Ndc80. (D) Microtubule decoration by Ndc80 and Cdk1-treated and untreated Ska. Scale bar: 5 µm.

#### Supplementary Figure 2.

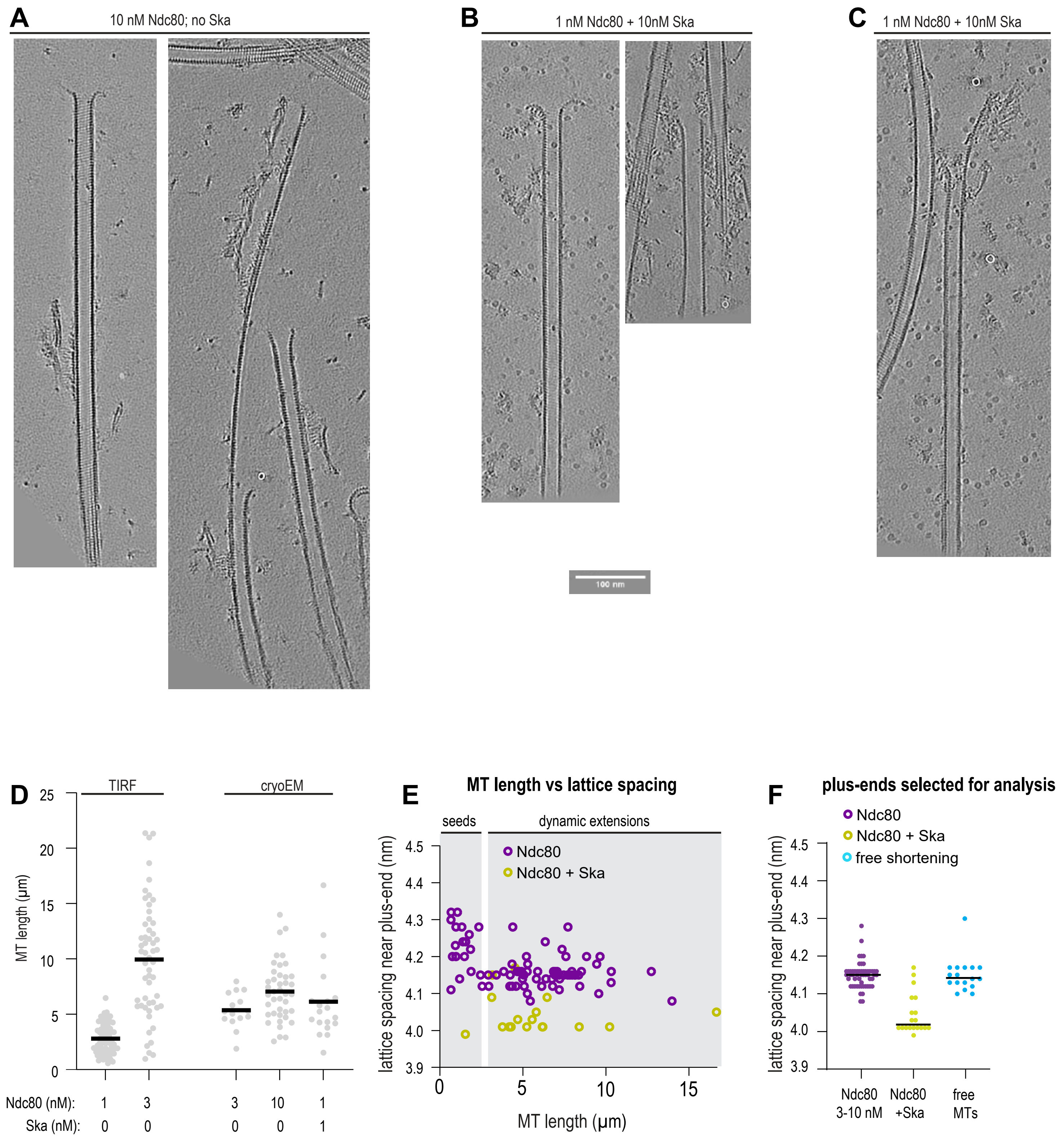


Additional examples of microtubules decorated with Ndc80 trains at 10 nM Ndc80 (A), with non-Ndc80 oligomers in presence of 1 nM Ndc80 and 1 nM Ska (B), and with Ndc80 trains stabilising extended sheet-like protofilaments in presence of 1 nM Ndc80 and 1 nM Ska (C). (D) Microtubule length determined using TIRF microscopy, or low-magnification cryoEM in conditions used to detect target positions for tomography, in presence of Ndc80 and Ska in indicated concentrations. Grey circles: individual microtubules, lines: median. (E) Correlation of tubulin lattice spacing and microtubule length in the samples containing Ndc80 only (magenta), or Ndc80 + Ska (yellow). (F) Lattice spacing near plus-ends of microtubules selected for further analysis of protofilament shapes.

Supplementary Figure 3.

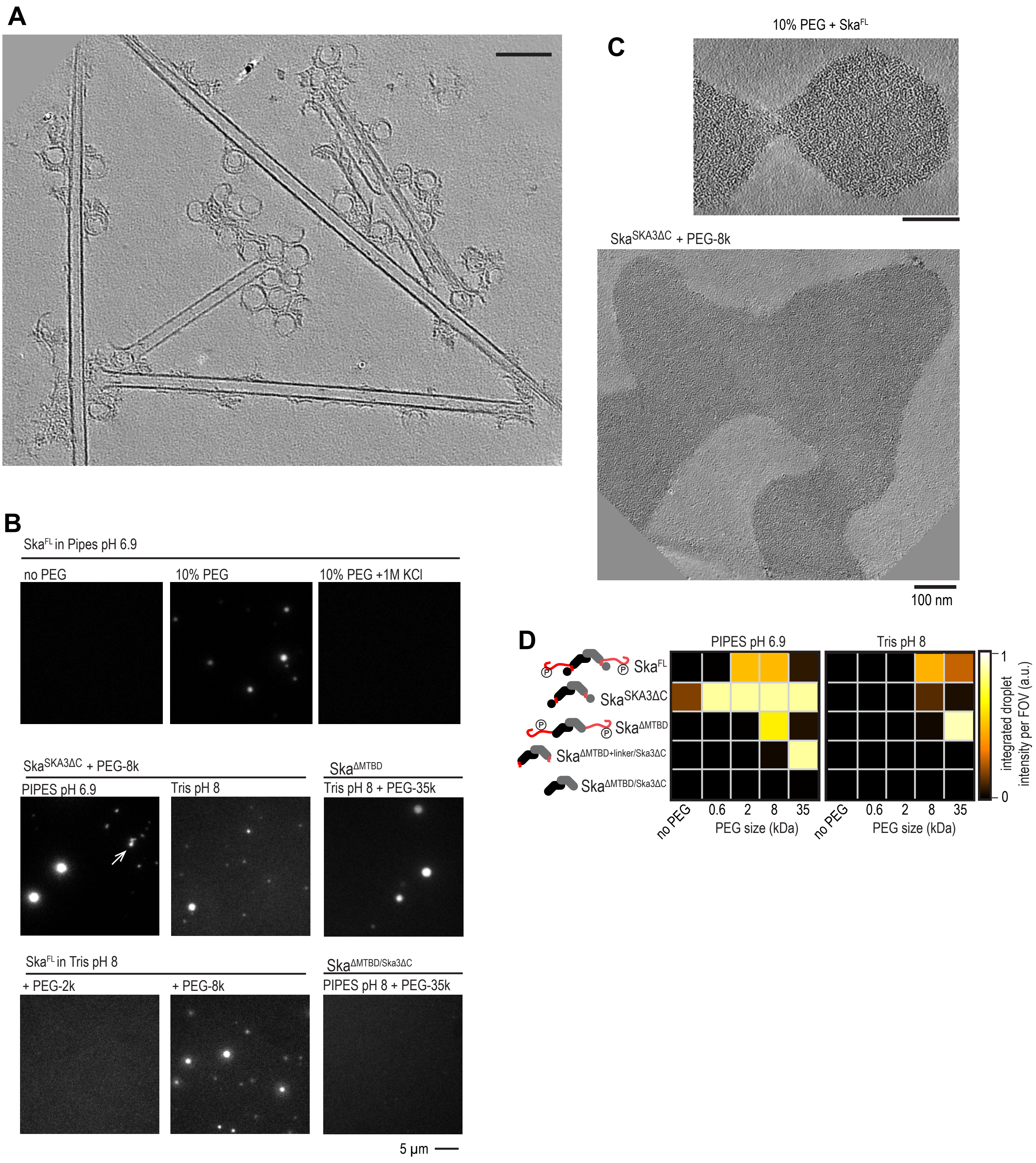


(A) A representative slice through a tomogram of a sample containing Ska and dynamic microtubules. (B) Fluorescence microscopy images of Ska in presence or absence of crowding agents and various buffer compositions indicated on the panel. Arrow points to non-spherical aggregates of SkaSKA3ΔC. (C) Slices through tomograms of FL Ska or SkaSKA3ΔC in presence of PEG. (D) Formation of fluorescent self-assembling oligomers of Ska in two various buffers, with the deletion construct indicated, and using a crowding agent indicated.

### Supplementary Figure 4.

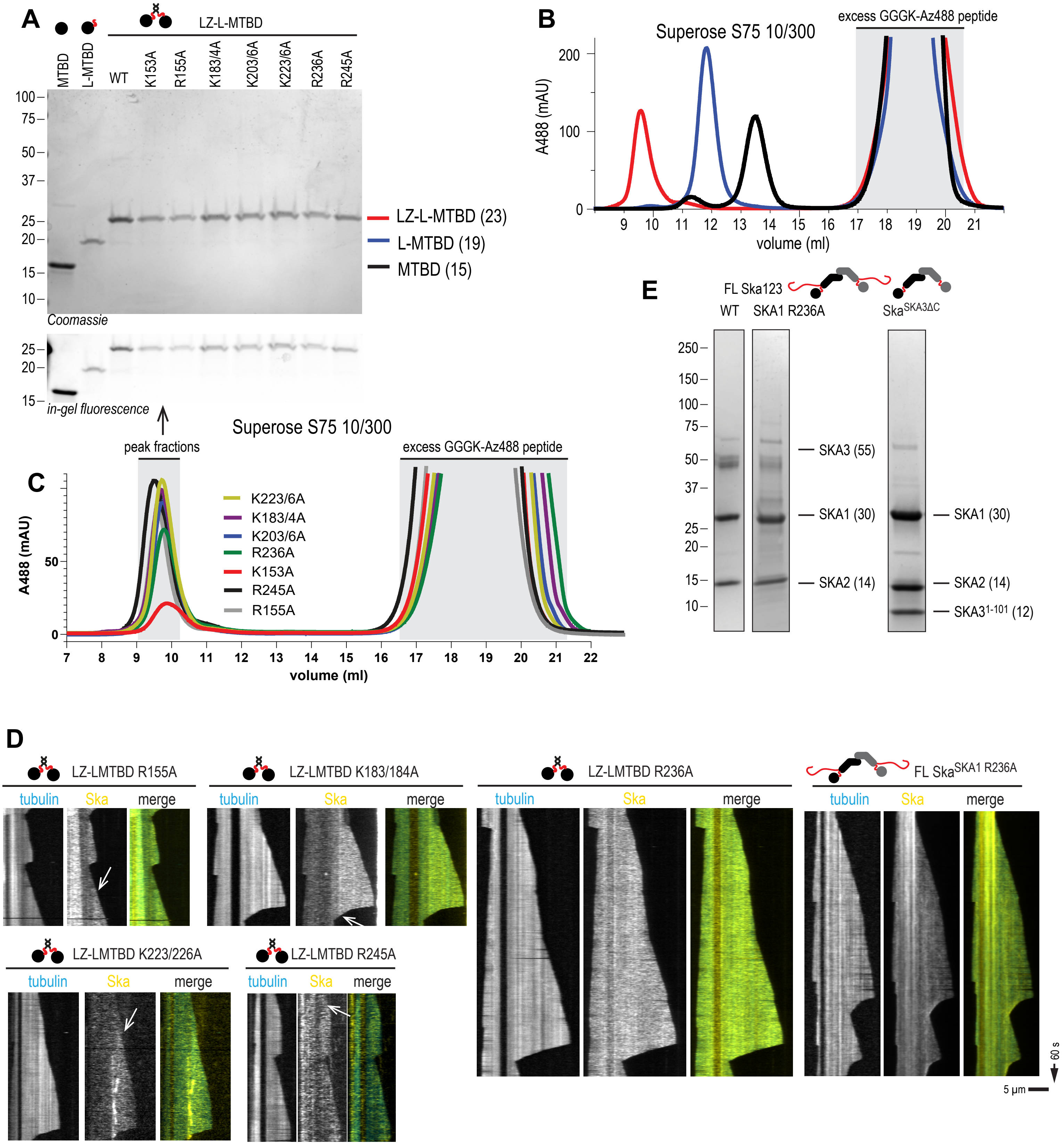


(A) SDS-PAGE of SKA1 MTBD constructs used in the study. (B) SEC profiles of dimeric LZ-LMTBD (red), and monomeric L-MTBD (blue) and MTBD (black) following fluorescent labelling with sortase. (C) SEC profiles of all point mutants of LZ-LMTBD following fluorescent labelling with sortase. (D) Examples of envelope formation assays for the LZ-LMTBD or FL Ska constructs indicated. Arrows point to a boundary between a brighter envelope and the rest of the lower intensity microtubule decoration.

#### Supplementary Figure 5.

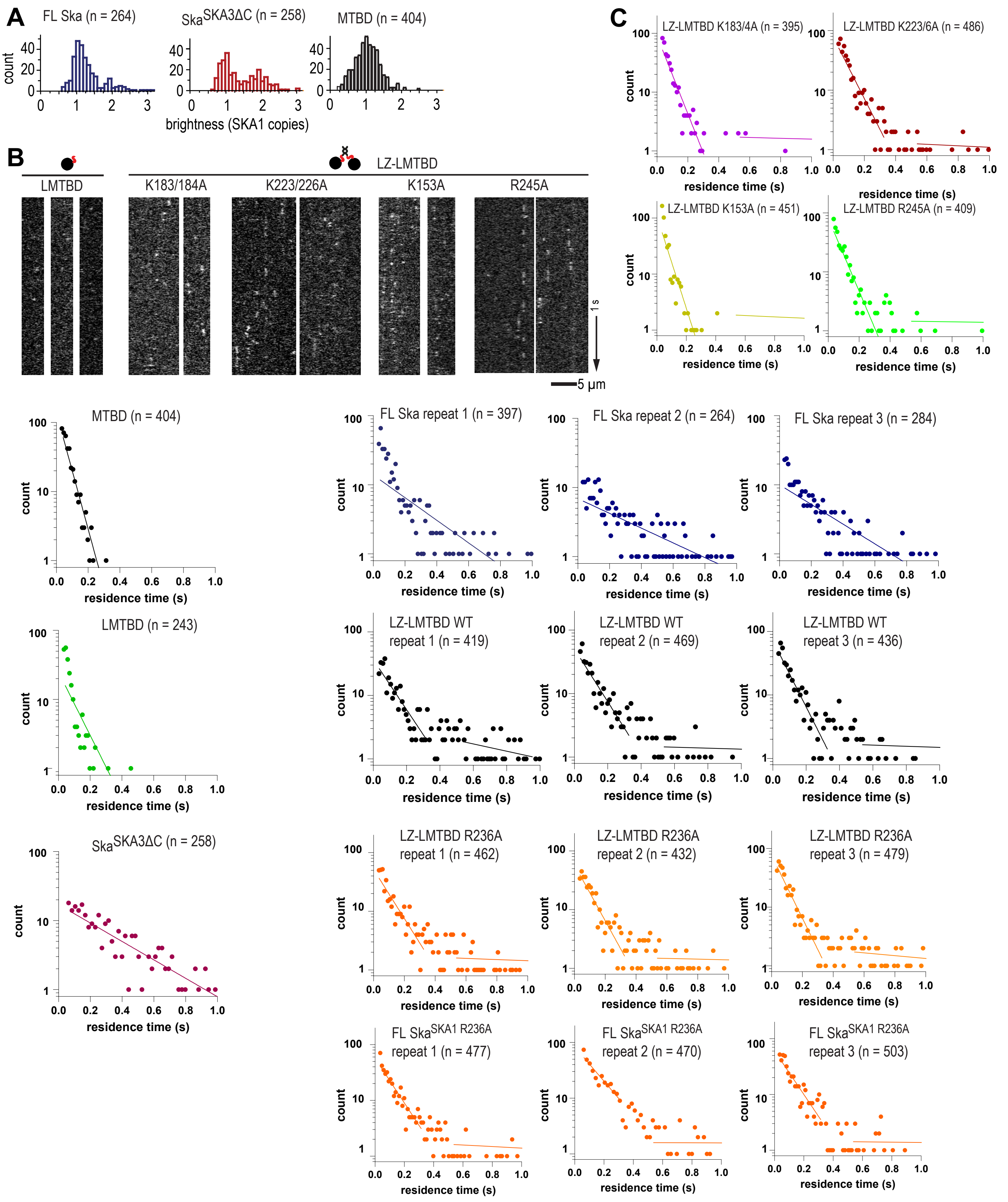


(A) Distribution of individual molecule brightness for the construct indicated. (B) Example kymographs used to determine single-molecule residence time of the construct indicated. (C) Distribution of residence times of the constructs indicated on each panel.

### Supplementary Figure 6.

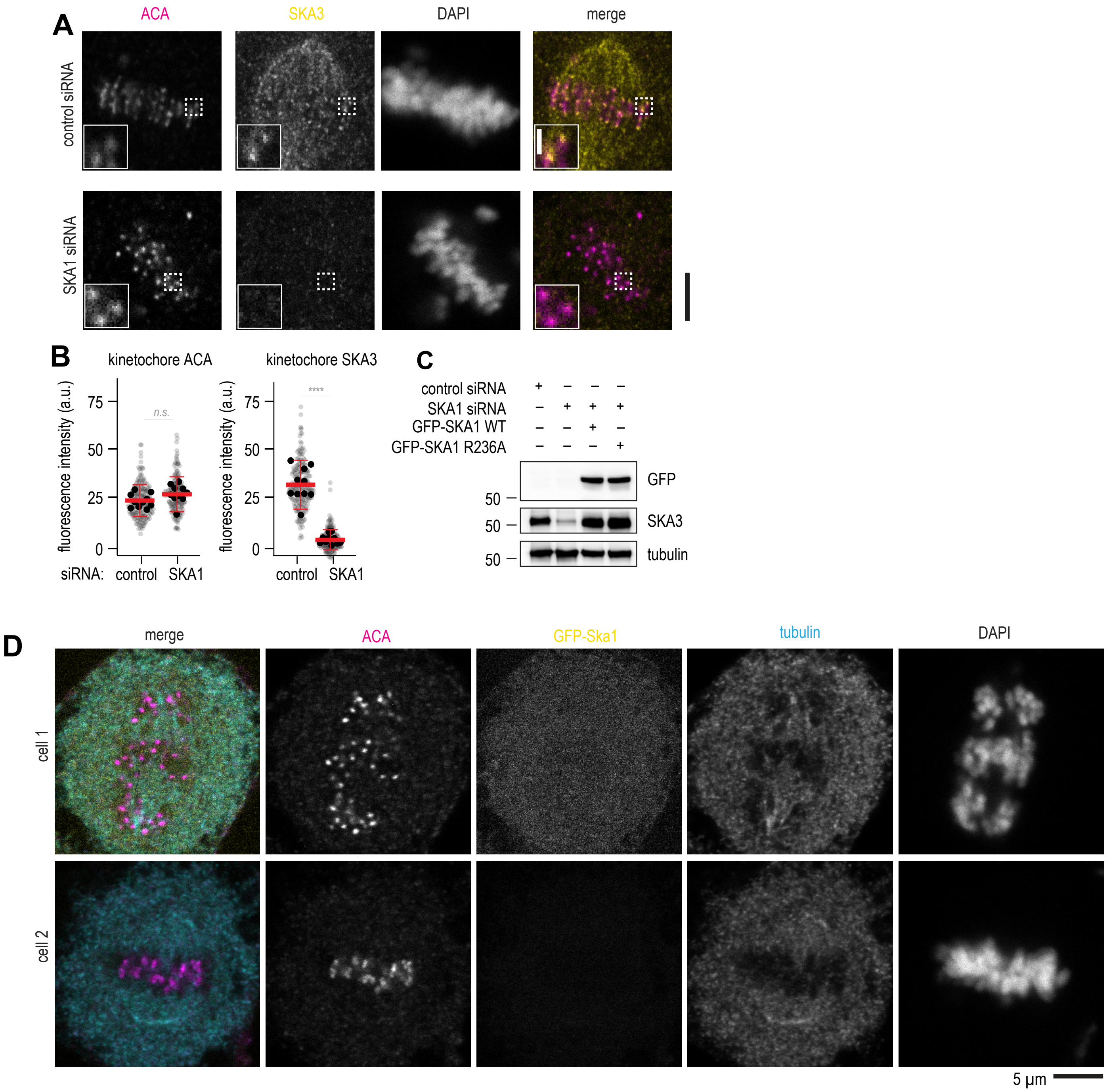


(A) Single planes from z-stacks of confocal images of cells treated with control siRNA or SKA1 siRNA and stained for ACA (magenta), SKA3 (yellow), and DAPI . (B) Quantification of SKA3 fluorescence intensity at kinetochores following treatment with the indicated siRNA. N = at least 20 kinetochores per cell (small symbols), 10 cells per condition (large symbols). Welch's t-test p value: ACA (control vs SKA1 siRNA): 0.0652; SKA3 (control vs SKA1 siRNA): < 0.0001. Lines show mean and S.D. (C) Western blots probing for GFP, SKA3, and tubulin as a loading control, using cells treated according to conditions indicated at the top of the panel. (D) Representative immunofluorescence confocal images of cold-treated cells with poor expression of GFP-Ska1, following siRNA of endogenous SKA1.
